# Objective Quality Assessment for Precision Functional MRI Data

**DOI:** 10.64898/2026.02.10.704857

**Authors:** Charles J. Lynch, Megan Chang, Immanuel Elbau, Evan M. Gordon, Timothy O. Laumann, Jingnan Du, Zach Ladwig, Maximilian Lueckel, Diana C. Perez, Indira Summerville, Jolene Chou, Megan Johnson, Claire Ho, Nicola Manfredi, Parsa Nilchian, Nili Solomonov, Eric Goldwaser, Tommy Ng, Stefano Moia, Cesar Caballero-Gaudes, Jonathan Downar, Fidel Vila-Rodriguez, Elizabeth Gregory, Zafiris J. Daskalakis, Daniel M. Blumberger, Kendrick Kay, Derrick Buchanan, Nolan Williams, Mahendra T. Bhati, Jacqueline Clauss, Benjamin Zebley, Lindsay W. Victoria, Jonathan D. Power, Logan Grosenick, Faith M. Gunning, Conor Liston

**Author notes:** Corresponding Authors: Charles J. Lynch and Conor Liston.

## Abstract

Precision functional mapping (PFM) enables individual-level characterization of brain network organization but requires substantially more and higher-quality fMRI data than is standard. Despite its growing use, objective criteria for data sufficiency and quality needed to ensure interpretable and replicable individual-level results remain unclear. Here, we introduce the Network Similarity Index (NSI), an objective measure of the extent to which functional connectivity (FC) patterns in an individual dataset express the large-scale network structure required for PFM. NSI captures the integrity of low-spatial-frequency, coherent network organization and denoising fidelity, and aligns closely with blinded expert assessments of PFM usability. NSI also accounts for variability in the rate at which FC becomes reliable across individuals. Here, we provide an open-source framework for NSI-based data quality evaluation and models for linking NSI values with expert-judged PFM suitability. This framework can also inform expected returns from additional data collection, enabling principled decisions about data sufficiency and replication in precision fMRI research.

## INTRODUCTION

Functional connectivity analysis of fMRI data, which assesses correlations in BOLD activity patterns across distributed brain regions ^1^, has provided a powerful means to non-invasively define large-scale functional brain areas and networks in humans ^2^. Early work established these functional networks primarily at the group level ^3,4^, by averaging relatively short duration fMRI runs across large cohorts to reveal a robust, canonical network architecture ^5,6^. These same connectivity principles are now applied within highly-sampled subjects to resolve inter-individual variation in the size, shape, and spatial boundaries of functional areas and networks^7–11^. This approach originated with the MyConnectome Project ^12,13^, which demonstrated that extensive longitudinal sampling of a single individual could reveal stable, individualized areal and network features ^14,15^. Over the past decade, this individual-level extension—termed precision functional mapping (PFM)^15,16^—has accelerated our understanding of human brain network organization ^11,13,14^, enabling new studies of individual differences in cognition ^10^ and psychopathology ^17–19^ while also motivating increasingly precise, circuit-informed interventions in psychiatric and neurologic disorders ^19–21^.

Individual-level mapping via PFM has uncovered organizational principles that were mischaracterized or undetectable by traditional group-based approaches. A prominent example comes from the motor system, where individualized mapping has challenged the classic view of primary motor cortex as a smooth homunculus ^22^ by revealing alternating effector-specific regions and inter-effector zones linked to action-control networks ^23,24^ that appear weakly in group averages but are sharply resolved within individuals. Similar principles extend beyond the motor cortex, with individual-specific network organizations spanning cortical zones ^25,26^, subcortex ^27–29^, and cerebellum ^7,30^ likewise obscured by group averaging ^11,31–33^.

Building on this foundation, PFM is now beginning to be applied in clinical populations ^34^, revealing atypical network features in neuropsychiatric disorders—for example, a nearly two-fold cortical expansion of the fronto-striatal salience network in individuals with major depression ^17,19,35^. In parallel, PFM has also identified pathophysiologically relevant network hyperconnectivity and treatment-responsive circuits in Parkinson’s disease ^34^. Moreover, by defining brain areas and circuits precisely within individuals, PFM has also enabled new dense-sampling longitudinal designs that track within-person fluctuations in circuit connectivity alongside symptoms ^17^ and endogenous ^36,37^ or experimentally induced physiological change ^38,39^ over time.

These advances have generated widespread enthusiasm for applying PFM more broadly. However, most PFM studies to date have adopted a “collect as much data as possible” approach, generating exceptionally large, high-quality fMRI datasets. As a result, new adopters are often left uncertain about whether their data are sufficient to support reliable individual-level network mapping, both when evaluating existing datasets and when designing new studies. In practice, this uncertainty has been addressed using indirect heuristics rather than by directly examining the data. Chief among these heuristics are total scan duration (i.e., “collect at least 30-40 minutes of data”), used as a proxy for sampling variability ^40^ and reliability ^14,16,41^, and head motion ^42^, used as an indicator of artifact burden. However, these heuristics may not always generalize: smaller datasets may suffice when signals of interest are clearly expressed, whereas larger datasets may fail when artifacts persist or relevant signals are attenuated. Consistent with this, our prior work ^43^ showed that multi-echo fMRI^44^—which improves BOLD contrast^45^ and removal of non-BOLD artifacts^46,47^—yields more reliable FC estimates than comparable or even larger amounts of legacy single-echo data, indicating that PFM suitability may depend as much on signal quality as on total scan duration.

Together, these limitations underscore the absence of a standardized approach for directly assessing dataset suitability for PFM based on the properties of the data itself. Currently, PFM suitability is often judged qualitatively through expert visual inspection of FC patterns, based on whether large-scale network structure is coherently expressed. When such expertise is unavailable—or when legacy datasets are repurposed without careful evaluation—PFM may be applied to data ill-suited for individualized mapping, increasing the risk of obscuring true individual-specific organization or mistaking artifactual patterns for novel network structure.

These errors have the potential for meaningful clinical consequences. For example, in circuit-targeted brain stimulation interventions, individualized maps are increasingly used both prospectively to precisely engage target circuits ^21^ and retrospectively to interpret network-level effects ^48^ and variability in treatment response when interventions succeed or fail ^49^. If maps are derived from datasets ill-suited for individualized inference, targeting errors may occur, and apparent variability in treatment response may reflect data limitations rather than true differences in target circuit engagement.

To address this problem, we developed the Network Similarity Index (NSI) as an objective measure of a dataset’s suitability for PFM. NSI is motivated in part by the observation that, in high-quality individual datasets, FC patterns reflect signal content at multiple spatial scales, including both a large-scale network structure that is broadly shared across individuals, as well as fine-scale and stable individual-specific network features. NSI quantifies the extent to which FC patterns express canonical network structure, thereby assessing whether large-scale organization is sufficiently resolved to support interpretation of individual-specific features. In doing so, NSI formalizes the qualitative process by which experts judge fMRI dataset quality into a reproducible quantitative framework.

We validate NSI across a broad, methodologically heterogeneous collection of datasets, including hundreds of patients with treatment-resistant depression, that reflect real-world artifact burden and clinical variability. Using these data, we show that NSI captures systematic variation in the integrity of large-scale FC organization, aligns with blinded expert judgments of dataset quality, and captures between-subject differences in the rate at which FC estimates become reliable with additional data. Here, we provide an open-source codebase for NSI-based quality evaluation and precision fMRI analysis, together with calibrated probabilistic models that map NSI values to expert-judged PFM usability, estimate the reliability of existing data, and inform expected returns from additional data collection. Together, these tools enable interpretable and generalizable decisions about dataset screening and data sufficiency for precision fMRI studies, spanning retrospective analyses and prospective protocol development.

## RESULTS

### Visual Inspection of Functional Connectivity for Large-Scale Network Organization

We begin by making explicit the qualitative rationale underlying the proposed NSI framework. In our laboratory and others, assessments of fMRI dataset suitability for PFM rely on visual inspection of FC maps, guided by expert judgment rather than formal criteria. This approach centers on evaluating whether FC maps exhibit coherent, large-scale spatial organization similar to known functional systems ^3,4^, thereby providing a shared organizational scaffold within which individual-specific features can be meaningfully interpreted.

In our workflow, FC maps are generated and inspected using *wb_view*, the graphical user interface for Connectome Workbench ^50^, with “dense dynamic” CIFTI files that allow rapid visual assessment of FC structure across the cortical surface and subcortical structures. By sampling FC at different brain regions, this inspection probes whether FC is dominated by spatially smooth, low–spatial-frequency structure that recapitulates canonical network motifs, or instead by fine-grained, spatially fragmented patterns that obscure network identity. **Figure 1** illustrates the qualitative contrast this inspection process is designed to detect using a single seed. Importantly, datasets with acceptable FC quality typically exhibit a coherent expression of large-scale network structure broadly across the brain, whereas datasets with poor FC quality tend to lack such structure nearly everywhere. Example movies of “crawling” FC seed maps, which systematically sample hundreds of seed locations, further illustrate this qualitative difference between high- and low-quality datasets (**Supplementary Materials Videos S1–2**).

**Figure 1.**
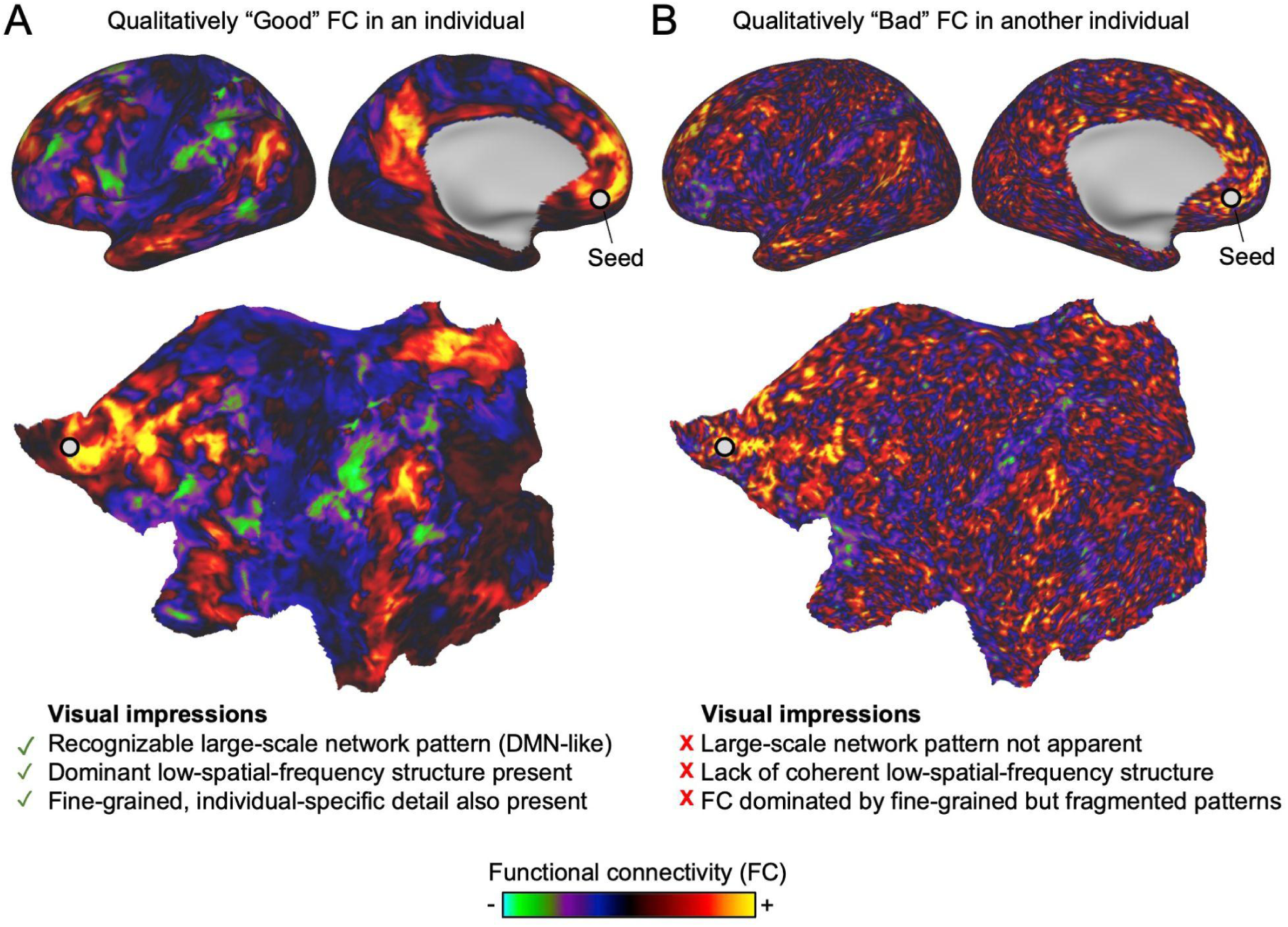
Visual inspection of seed-based functional connectivity (FC) maps to assess PFM suitability. (**A**) Qualitatively “good” FC: in one representative individual dataset, a seed placed in ventral medial prefrontal cortex (vmPFC) yields a spatially smooth, low–spatial-frequency FC pattern with contiguous positive and negative structure that resembles a canonical large-scale network motif (DMN-like). Fine-grained structure is also evident but expressed within this coherent large-scale organization. (**B**) Qualitatively “bad” FC: in a different individual dataset, the same seed yields an FC map dominated by fine-grained, spatially fragmented structure with rapid sign reversals and intermingled positive and negative values across neighboring vertices, obscuring recognizable large-scale network organization. Color scales depict FC strength (warm = positive; cool = negative). Bottom panels show the same FC maps projected onto two-dimensional cortical flatmaps, and the checklists summarize the qualitative visual criteria commonly used during manual QC to assess the presence or absence of coherent large-scale network organization. No overt spatial smoothing was applied to either dataset displayed here.

In higher-quality datasets, a seed placed at nearly any location yields FC patterns with broad, contiguous structure that varies smoothly across space. These low–spatial-frequency patterns resemble familiar large-scale brain networks and provide a shared organizational backdrop across individuals. **Fig. 1A** shows one such example, in which a seed placed in ventral medial prefrontal cortex (vmPFC) produces an FC map that resembles a default mode network–like motif. The same seed in a lower quality dataset (**Fig. 1B**) instead yields FC maps dominated by fine-grained, spatially irregular structure, characterized by rapid sign reversals and intermingled positive and negative values across neighboring vertices. These kinds of maps lack recognizable large-scale structure, making it difficult to identify known networks or to interpret finer-scale features as meaningful individual variation.

The visual impressions summarized in **Fig. 1** reflect a shared working intuition in the field regarding individual-level FC measurements. Specifically, FC is organized around a recognizable, large-scale spatial structure that is shared across individuals, providing a common organizational framework within which finer-grained, individual-specific features—and systematic deviations from that shared structure—can be meaningfully expressed, interpreted, and compared across people.

### Functional Connectivity Contains Shared Large-Scale and Individual-Specific Components

The descriptive observations of FC structure described above point to a potential organizing principle of FC across spatial scales. In particular, FC maps that are widely regarded as suitable for PFM often appear to be dominated by a low–spatial-frequency component that is similar across individuals, giving rise to recognizable large-scale network structures that can be aligned across people ^51^. At the same time, experienced investigators note that FC maps derived from high-quality data also contain additional fine-scale structure that is stable within individuals ^11,14^ and gives rise to reliable, person-specific features expressed relative to this shared large-scale organization. Although these qualitative impressions suggest a separation between shared large-scale structure and individual-specific fine-scale organization, the two components have not been explicitly isolated or quantitatively evaluated.

To formalize the intuition that FC comprises a conserved large-scale (low–spatial-frequency) scaffold upon which stable, individual-specific fine-scale structure is expressed, we analyzed the spatial frequency composition of FC in a cohort of n = 31 highly sampled individuals. Each individual contributed more than 240 minutes of motion-censored resting-state fMRI data, enabling within-individual split-half reliability analyses. FC maps were computed for many seed locations distributed evenly across the cortical surface and decomposed into a low–spatial-frequency component and an orthogonal fine-scale component using a surface-based spatial frequency decomposition. **Figure 2A** illustrates this decomposition of the DMN-like FC pattern from **Fig. 1A** into smooth low–spatial-frequency and fine-scale components, shown separately for split halves of the data. In all n = 31 highly-sampled individuals, FC variance was concentrated in low spatial frequencies (70.3 ± 6.2%), corresponding to coarse, smoothly varying patterns that span large portions of cortex (**Fig. 2B**). Both components appeared similar across the split halves in the example individual (**Fig. 2A**). To test this across the full sample, we performed a spatial similarity analysis to quantify within-individual reliability, between-individual similarity, and correspondence with network FC templates, providing an additional test of cross-subject generalization.

**Figure 2.**
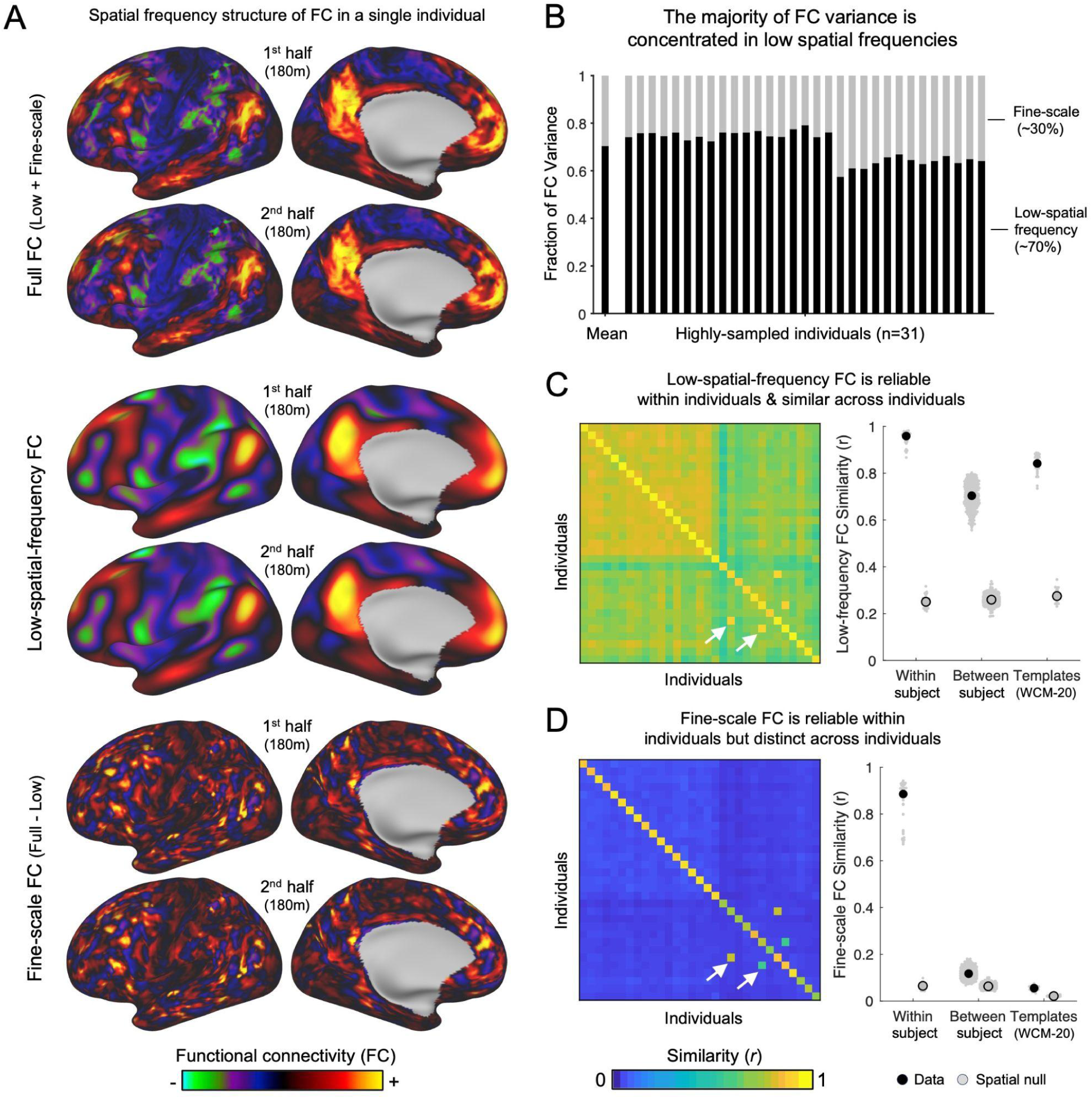
Functional connectivity comprises a conserved low–spatial-frequency scaffold and individual-specific fine-scale structure. **(A)** Seed-based functional connectivity (FC) maps from a single highly sampled individual (ME01 ^43^; ∼6 hours total of resting-state fMRI data, split into two independent halves of approximately 3 hours each). Maps are shown for the full FC estimate (top), its low–spatial-frequency component (middle), and the residual fine-scale structure (bottom) obtained by subtracting the low-frequency component from the full map. Each component is displayed separately for the first and second halves of the data (180 minutes per split in this individual). Both low-frequency and fine-scale components exhibit high split-half reliability within the individual, despite qualitatively distinct spatial organization. **(B)** Across individuals, the majority of FC variance is concentrated in low spatial frequencies. Bars show the fraction of FC variance attributable to low–spatial-frequency structure (black) and residual fine-scale structure (gray) for each individual (n = 31), with the group mean shown at left. **(C)** Low–spatial-frequency FC is reliable within individuals and similar across individuals. Individual x individual similarity matrices quantify within-individual split-half reliability (diagonal) and between-individual similarity (off-diagonal) for low–spatial-frequency FC (n = 31, each contributing >240 minutes of motion-censored fMRI data). Summary plots contrast within-individual and between-individual similarity values, alongside correspondence with group-level network FC templates (WCM-20 ^17^). Low–spatial-frequency FC shows strong within-individual reliability, substantial cross-individual similarity, and high correspondence with group templates, indicating a conserved large-scale scaffold that is consistently expressed across individuals and datasets. White arrows highlight two individuals (MSC02, MSC06) from the Midnight Scan Club ^16^ who also participated in the Cast-Induced Plasticity ^38^ study (CAST01, CAST02). These repeat individuals provide an internal positive control on the similarity metric: across-study data from the same person show elevated similarity relative to between-individual comparisons. To avoid inflating within-individual summary statistics, these individuals are excluded from within- versus between-individual aggregates but retained in the full similarity matrices to illustrate cross-dataset within-individual correspondence. **(D)** In contrast, residual fine-scale FC structure is reliable within individuals but distinct across individuals. Similarity matrices and summary plots show high within-individual split-half reliability but low between-individual similarity and minimal correspondence with group-level FC templates.

Low–spatial-frequency FC showed high split-half reliability within individuals (median *r* = 0.96 ± 0.03), along with strong similarity both across individuals (median *r* = 0.70 ± 0.05) and to the network FC templates (median *r* = 0.86 ± 0.04). These patterns are evident in the similarity matrix as strong diagonal and off-diagonal structure (**Fig. 2C**) and exceeded those from a spatial null model by a wide margin (all z-scores > 20), indicating a conserved, shared component of functional organization not explained by generic spatial smoothness. In contrast, residual fine-scale FC showed high split-half reliability within individuals (median *r* = 0.89 ± 0.03) but weak similarity across individuals (median *r* = 0.12 ± 0.02) and to network FC templates (median *r* = 0.06 ± 0.001; **Fig. 2D**). Although these values exceeded those expected under the spatial null (all z-scores > 5), their absolute magnitudes were small compared with the much larger similarities observed for low–spatial-frequency FC (*r* ≈ 0.7–0.9). Overall, this pattern indicates that fine-scale FC organization is stable within individuals yet largely idiosyncratic across individuals, with minimal shared structure at the group level.

Together, the results in **Fig. 2** establish a consistent empirical relationship between the resolution of shared low–spatial-frequency FC structure and the reliable recovery of individual-specific FC. Across individuals, low-frequency FC reflects a conserved organizational scaffold, while fine-scale residual FC captures stable, person-specific deviations from this shared reference. Although individual specificity resides primarily in fine-scale structure, noise degrades both low- and high-frequency components. Crucially, only the low–spatial-frequency component can be evaluated against an external reference, because its expected large-scale organization is known a priori. As a result, the presence of a well-resolved low-frequency scaffold is an indicator that the data are of sufficient quality to support interpretable individual-level FC estimates. In the following section, we introduce an objective measure that quantifies how well low–spatial-frequency FC structure is resolved, providing an empirical basis for determining whether a given dataset is suitable for PFM.

### The Network Similarity Index (NSI) for PFM

To formalize the manual inspection procedures used to judge whether a dataset is suitable for PFM, we developed the Network Similarity Index (NSI). NSI formalizes the criterion implicitly used during manual data inspection—namely, whether seed-based FC maps exhibit coherent large-scale network organization rather than a noise-dominated spatial structure.

Specifically, NSI assesses how well seed-based FC maps can be approximated by weighted combinations of canonical functional brain networks (**Fig. 3**). For each seed location, the observed FC map is reconstructed using ridge regression with a set of user-specified network templates (see **Supplementary Fig. 1**). This procedure yields a predicted FC map and an associated set of regression weights, which are shown for illustrative purposes to demonstrate the network composition of the reconstruction for an example seed (**Fig. 3A–B**). Importantly, the primary quantity of interest is the goodness-of-fit between the observed and reconstructed FC patterns. This goodness-of-fit, quantified as variance explained (R²), defines the NSI value for that seed (**Fig. 3C**) and serves as the sole metric used for subsequent quality assessment analyses.

**Figure 3.**
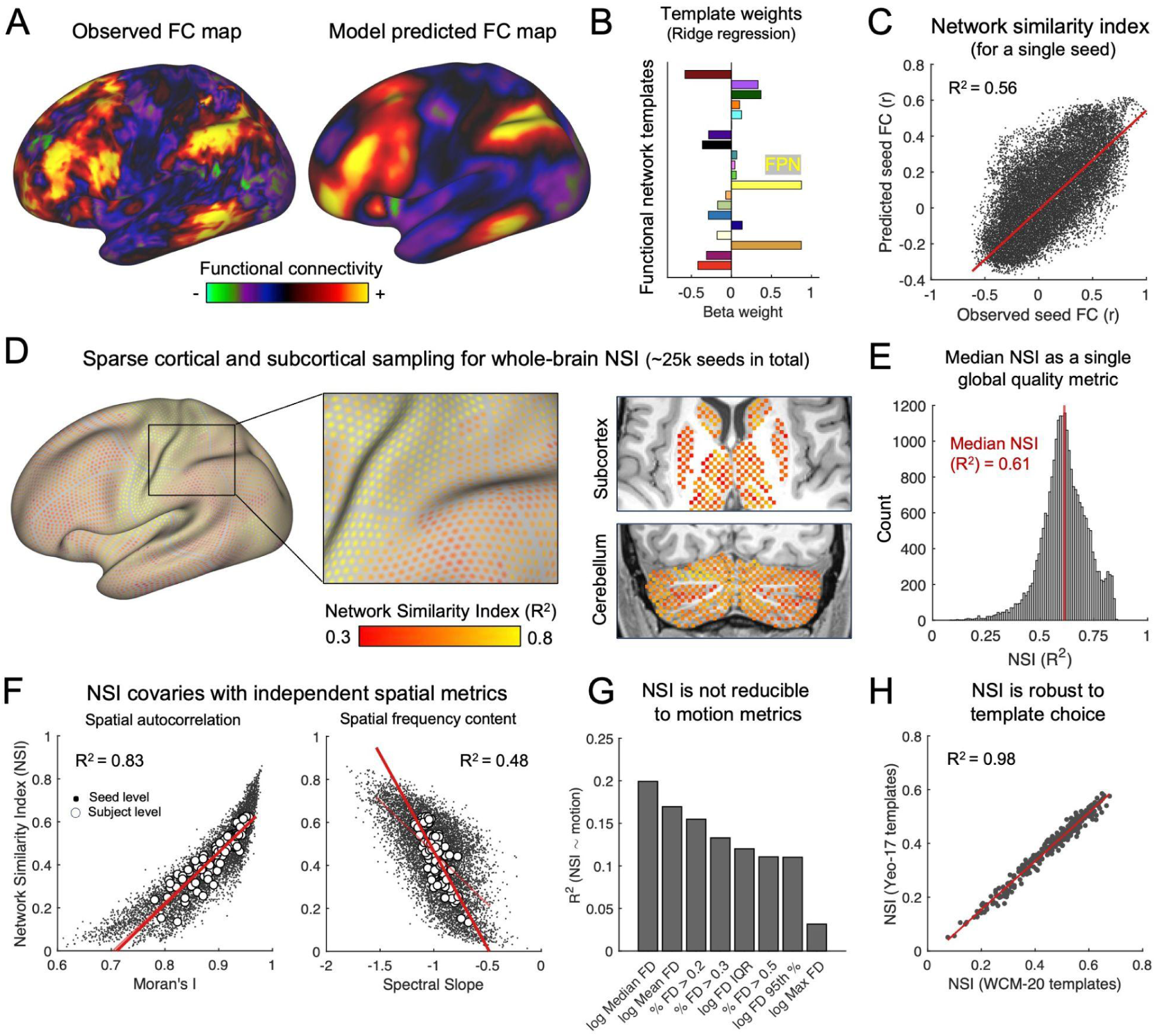
Overview of the Network Similarity Index (NSI) and its relationship to large-scale spatial organization. NSI provides an algorithmic analog of expert manual quality control by quantifying the extent to which seed-based functional connectivity (FC) patterns express a coherent, shared low–spatial-frequency network scaffold. Specifically, NSI measures how well individual FC maps can be reconstructed as weighted combinations of canonical functional network templates, reflecting the presence of recognizable large-scale network organization. **(A)** Observed seed-based FC map for a single cortical seed (left) and the corresponding FC map predicted by ridge regression using canonical network templates (right). **(B)** Ridge regression weights (β) for the same seed, showing the relative contributions of individual network templates to the reconstructed FC map. **(C)** Scatter plot of observed versus predicted FC values for all vertices for the same seed; the linear fit (red) quantifies variance explained (R²), which defines the single-seed NSI. **(D)** Sparse cortical, subcortical, and cerebellar sampling scheme used to compute whole-brain NSI (∼25,000 seeds total) to provide computationally tractable whole-brain coverage. Insets illustrate spatial coverage and the resulting seed-wise NSI values mapped back to anatomy. **(E)** Distribution of seed-wise NSI values for a single dataset; the median NSI value serves as a robust global summary measure. **(F)** NSI covaries with independent spatial metrics indexing the integrity of the shared low–spatial-frequency scaffold. NSI increases with greater spatial autocorrelation (Moran’s I; left) and decreases with higher spatial frequency content (spectral slope; right), consistent with coherent large-scale organization. Small black points denote single-seed values; larger white points denote subject-level averages. Red lines indicate linear fits, with R² summarizing association strength. **(G)** NSI shows only modest associations with multiple head-motion summaries (all log-transformed). **(H)** Subject-level median NSI values computed using two different canonical network template sets (WCM-20 ^17^ vs. Yeo-17 ^4^) are highly concordant, demonstrating that NSI estimates are robust to network FC template choice. Panels **A–E** show illustrative examples from a single highly-sampled healthy individual (“ME01” ^43^), selected to demonstrate NSI computation and visualization; all quantitative analyses in panels **F–H** are performed across the full validation cohort.

To provide whole-brain coverage while remaining computationally tractable, NSI is computed using a sparse but comprehensive sampling of cortical, subcortical, and cerebellar seed locations (**Fig. 3D**). The median of all within-individual seed-wise NSI values is then computed, representing a single global summary measure (**Fig. 3E**) that remains stable across a broad range of ridge regularization parameters (**Supplementary Fig. 2**). In this way, NSI provides an objective, algorithmic analog of the manual review procedure described in **Fig. 1**, while remaining computationally efficient and requiring only a few minutes per dataset on a standard workstation.

### Validating NSI as a Measure of Large-Scale FC Organization

To validate NSI as a measure of large-scale functional organization relevant for PFM, we evaluated (i) its relationship to independent spatial metrics that index the smooth, slowly varying spatial organization of functional connectivity, (ii) its correspondence with head-motion measures, and (iii) its sensitivity to the choice of canonical network templates. Analyses were conducted in a randomly sampled subset of individuals (n = 100) drawn from a heterogeneous corpus of >300 fMRI datasets (see **Supplementary Materials**).

Because NSI is designed to quantify how well an individual’s FC maps resolve the smooth, large-scale organization of functional networks that is shared across individuals, a natural validation strategy is to compare NSI against independent measures that index the spatial properties of network template FC maps themselves. We therefore focused on spatial metrics that index smooth, slowly varying FC organization, specifically spatial autocorrelation (Moran’s I) and the relative dominance of low–spatial-frequency spatial modes, each of which provides an independent assessment of the shared large-scale scaffold.

As expected, NSI increased with greater spatial autocorrelation (Moran’s I; R² = 0.83) and with greater dominance of low–spatial-frequency structure (i.e., a more negative spectral slope; R² = 0.48). These associations were quantified using seed-level linear mixed-effects models with subject as a random effect, capturing both within-subject (across seeds) and between-subject contributions to variance in NSI (**Fig. 3F**). Together, these relationships indicate that NSI is sensitive to coherent, smoothly varying FC patterns expressed over large spatial scales.

Given that head motion has traditionally served as a primary proxy for fMRI data quality through its association with artifact burden, we next examined the relationship between NSI and motion summary measures. NSI showed only modest associations with head-motion metrics, with no measure explaining more than 20% of NSI variance (**Fig. 3G**). This limited correspondence indicates that NSI is associated with—but not reducible to—conventional motion-based quality measures, instead reflecting properties of functional organization that persist beyond what is indexed by head motion alone.

A potential limitation of NSI is its reliance on a user-specified set of canonical network templates, raising the possibility that NSI estimates could vary with the choice of network definition. However, we found that NSI values were nearly identical when computed using different canonical network template sets (WCM-20 ^17^ vs. Yeo-17^4^; **Fig. 3H**), with subject-level median NSI values showing near-perfect correspondence across template definitions (R² = 0.98). This result indicates that inter-individual differences in NSI are preserved across these two network template sets, despite variation in how particular large-scale functional networks are defined.

### NSI Decreases with Progressive Removal of BOLD-Like Signals

Having established that NSI is sensitive to the integrity of large-scale, spatially coherent FC structure, we next tested whether it is sensitive to denoising fidelity. To do so, we systematically perturbed our multi-echo ICA (ME-ICA ^47,52^) denoising pipeline by progressively swapping components classified as signal with an equal number classified as noise. In n=100 adult participants, each contributing four 15-min multi-echo resting-state runs, we generated progressively degraded versions of the data by swapping components classified as signal with an equal number classified as noise (**Fig. 4A**). Seed-based FC from a representative individual illustrates the resulting degradation. At baseline, a posterior cingulate cortex (PCC) seed yields a spatially smooth FC map with a recognizable large-scale network motif; as the fraction of swapped components increases, this coherent structure is progressively disrupted, giving way to diffuse, poorly organized FC patterns (**Fig. 4B**).

**Figure 4.**
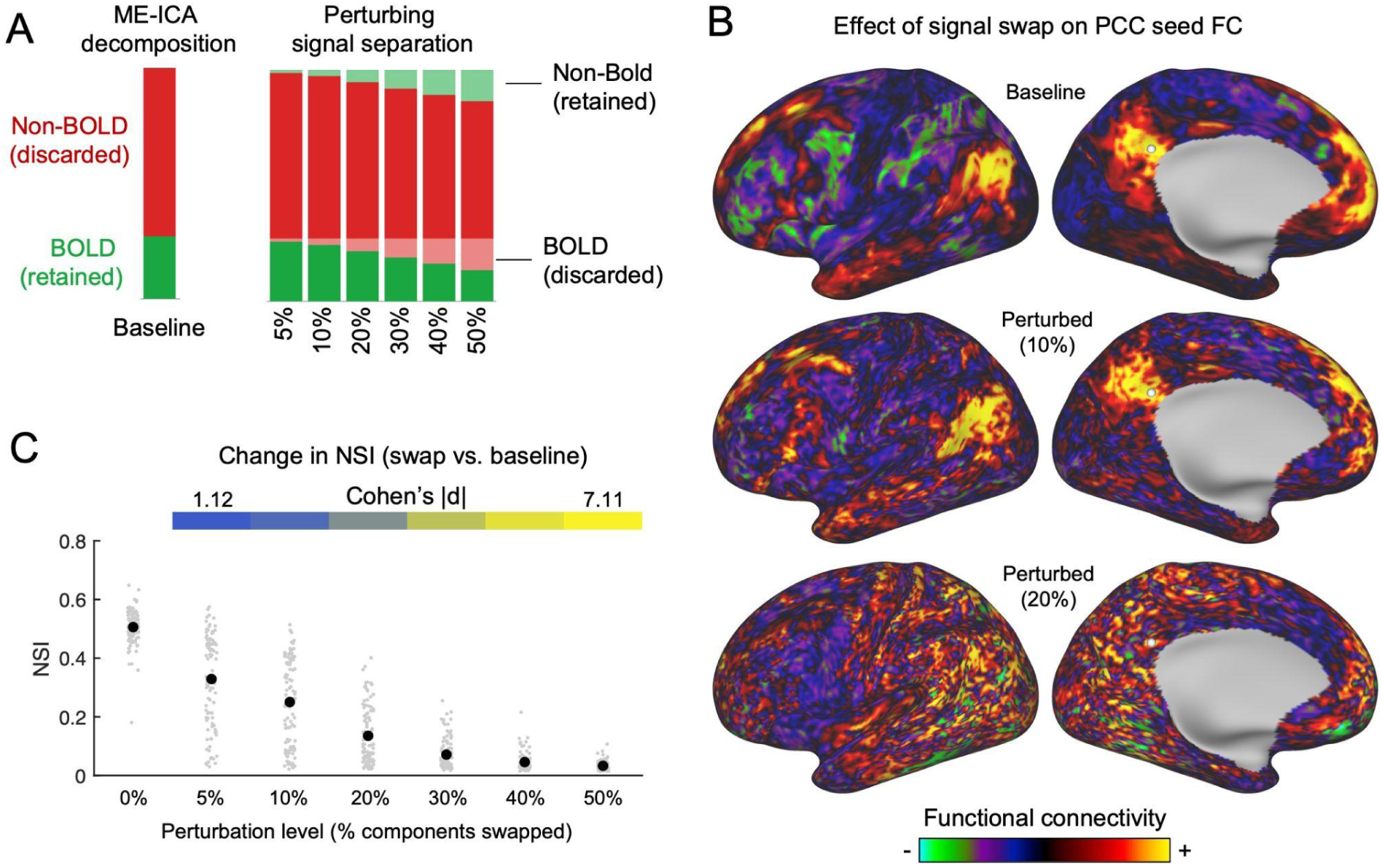
NSI tracks denoising fidelity and is most sensitive to loss of BOLD-like signal variance. **(A)** Schematic of the perturbation procedure. Starting from the baseline and ground truth ME-ICA decomposition, an increasing fraction of components labeled as BOLD-like were swapped with an equal number labeled as non-BOLD-like, generating a graded misclassification series (5%-50% swapped). **(B)** Effect of signal swapping on single-seed functional connectivity (FC) in a representative individual. At baseline, a seed in the posterior cingulate cortex (PCC) yields a spatially smooth, low-frequency FC pattern expressing a recognizable large-scale network motif. With increasing swap fractions, this coherent structure is progressively obscured by diffuse, spatially structured noise. **(C)** Group-level effects of signal swapping on spatial structure measurements across 100 individuals. Gray points show individual subjects; black points indicate subject means. Heat strips above each panel denote effect sizes (Cohen’s |d|) comparing each swap level to the baseline (0% swapped). NSI shows a graded, swap-fraction-dependent decline, with even mild perturbations producing large effects and higher swap fractions yielding large deviations from baseline.

Across subjects, NSI decreased monotonically with increasing swap fraction (**Fig. 4C**), indicating graded degradation of large-scale network structure as BOLD-like signals were removed. **Figure 4C** summarizes subject-level changes in NSI relative to each individual’s baseline, with gray points showing per-subject values and black markers indicating group means. NSI of perturbed data exhibited substantial decreases relative to baseline, with effect sizes that were large even for mild perturbations (|d| ≈ 1 at f05) and increased sharply with more aggressive swapping (|d| > 5 by f30), reflecting highly consistent disruption across individuals.

To determine whether these decreases primarily reflected the removal of BOLD-like signal variance or the introduction of non-BOLD variance, we modeled change in NSI as a function of the total variance explained by discarded signal components and introduced noise components, using mixed-effects analyses with subject as a random effect. Both factors independently reduced NSI, with greater decreases associated with removal of BOLD-like signal variance than with addition of non-BOLD variance. In a mixed-effects model predicting ΔNSI, the standardized coefficient for removed BOLD-like variance was more negative (β_RemoveSignal_ = −0.071, 95% CI [−0.082, −0.061]) than that for added non-BOLD variance (β_AddNoise_ = −0.048, 95% CI [−0.061, −0.034]). Accordingly, the difference between coefficients (β_RemoveSignal_ − β_AddNoise_ = −0.024, 95% CI [−0.046, −0.001]) was statistically significant (p = 0.039, Wald test), indicating a stronger impact of signal removal on NSI degradation. Together, these results indicate that NSI is sensitive to denoising fidelity, decreasing both when spatially structured BOLD signal is removed and when structured non-BOLD variance is introduced.

### NSI Tracks Expert Judgments of Dataset Quality

Manual assessment of fMRI dataset suitability for PFM is typically based on expert visual inspection of seed-based FC maps, guided by qualitative judgments about the presence or absence of coherent large-scale network organization (**Fig. 1**). To test whether NSI recapitulates these established expert judgments, we compared NSI values to blinded ratings of dataset quality provided by experienced investigators. Six independent raters (study authors ML, TOL, EMG, ZL, JD, and DCP) were recruited to evaluate 25 preprocessed, denoised, surface-registered resting-state fMRI datasets, with 5 datasets covertly duplicated to assess intra-rater reliability, yielding 30 total dataset evaluations.To ensure a broad range of data quality, the datasets selected intentionally spanned newer high-quality acquisitions and legacy datasets, enabling evaluation of NSI across the full spectrum of PFM usability. Raters were provided access to anonymized datasets and a standardized evaluation rubric. For each dataset, raters (i) inspected FC maps using wb_view (Connectome Workbench ^53^), (ii) assigned a continuous quality score from 1–10, and (iii) rendered a binary judgment indicating whether the dataset was usable for PFM (Y/N).

Blinded raters showed broad agreement in continuous quality ratings across datasets (mean pairwise inter-rater correlation r = 0.65 ± 0.16; **Fig. 5A**), with datasets ordered by NSI (top to bottom) and raters ordered by their correspondence with NSI (left to right). Majority agreement in binary usability decisions across raters exhibited a pronounced quadratic relationship with NSI (**Fig. 5B**; quadratic term R² = 0.454, p < 0.001; linear term R² = 0.009), with high agreement for datasets at low and high NSI values and reduced agreement at intermediate NSI levels. This pattern indicates that expert judgments converge more at the extremes of data quality but diverge more for borderline datasets, highlighting the inherent ambiguity of thresholded usability decisions and motivating the need for an objective, continuous quality measure such as NSI. Across the six blinded raters, NSI showed a consistent linear association with continuous quality ratings (Pearson r range across raters: 0.66–0.90; see **Supplementary Fig. 3** for individual-rater plots). Intra-rater reliability estimated from duplicated datasets was high overall (Pearson *r* range: 0.76–0.99), indicating good rater consistency. By contrast, associations between blinded ratings and total scan duration were weaker and more variable across raters (Pearson *r* range: 0.29–0.69), suggesting that data quantity alone does not fully account for perceived dataset quality.

**Figure 5.**
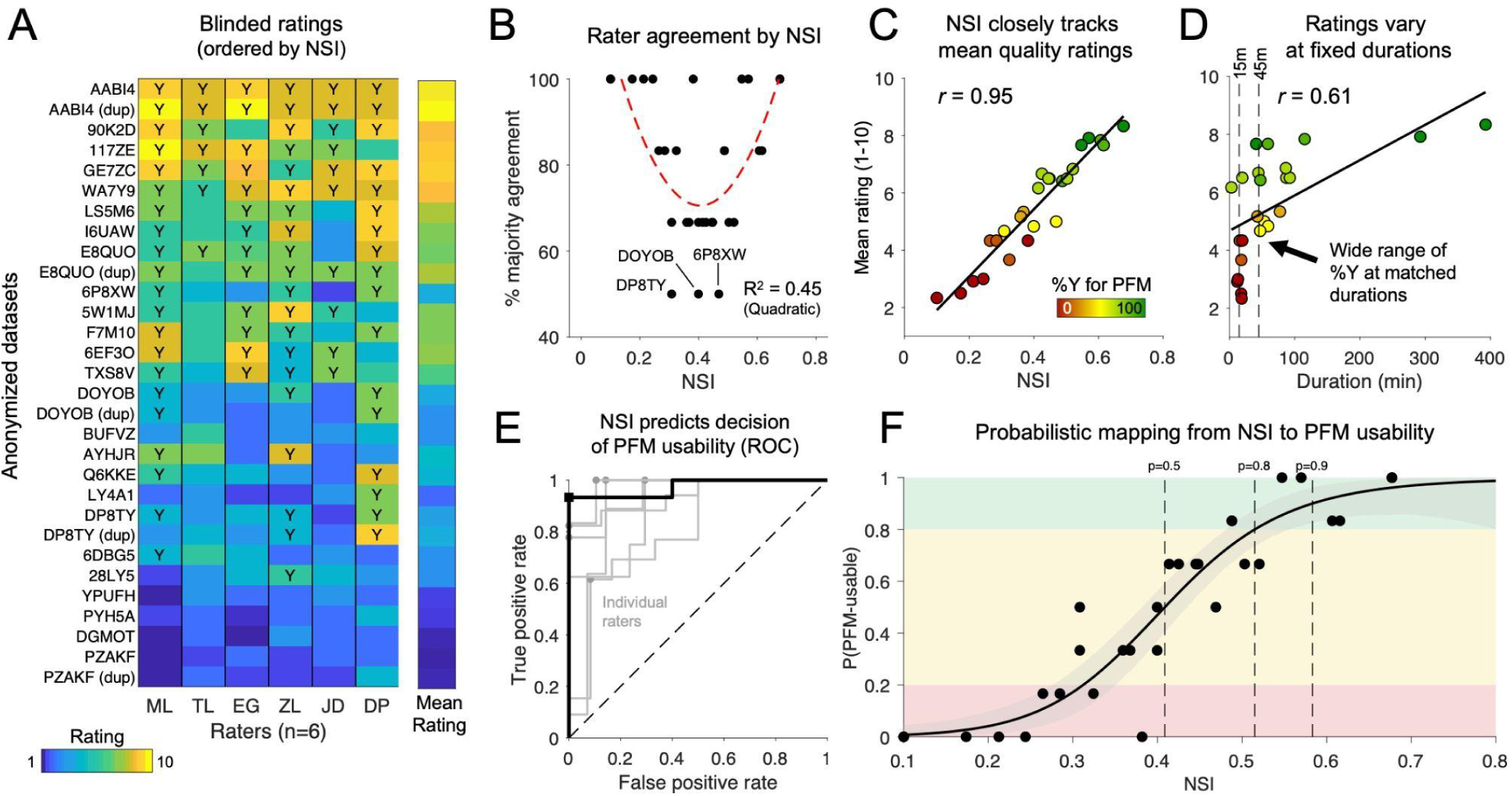
NSI predicts blinded assessments of PFM dataset usability and supports a calibrated decision framework. **(A)** Blinded quality ratings from six independent raters for anonymized fMRI datasets, ordered by Network Similarity Index (NSI). Heatmap colors indicate continuous quality scores (1–10), with overlaid “Y” denoting binary judgments that a dataset was usable for precision functional mapping (PFM). Raters show strong concordance in relative dataset ordering. *Dup* indicates a duplicate dataset (raters blinded). **(B)** Majority rater agreement (the fraction of six independent raters agreeing with the modal binary decision for each dataset) as a function of NSI, revealing lowest consensus at intermediate NSI values, consistent with ambiguity near the usability boundary. **(C)** Mean blinded rating versus NSI, demonstrating a strong monotonic relationship between NSI and perceived dataset quality. Point color indicates the proportion of raters endorsing PFM usability. **(D)** Mean blinded rating versus total data duration (minutes). Vertical dashed lines indicate example matched durations, highlighting wide variability in perceived quality for datasets with similar scan lengths. **(E)** Receiver operating characteristic (ROC) curves illustrate discrimination of PFM usability by NSI. Gray curves show individual raters, and the black curve shows the pooled model across raters. **(F)** Calibration curve mapping NSI to the probability that a dataset is judged PFM-usable by blinded raters. Points reflect per-dataset rater consensus; the solid line shows the fitted model with shaded bootstrap 95% confidence intervals. Background shading denotes low, intermediate, and high usability confidence regimes, and dashed vertical lines mark NSI values corresponding to 50%, 80%, and 90% predicted usability.

When ratings were aggregated across raters, higher mean blinded quality scores coincided with a greater fraction of raters endorsing PFM usability (%Y; **Fig. 5C**). NSI exhibited a near-monotonic relationship with both measures, progressing from low mean scores and few usability endorsements to high scores and near-unanimous agreement (*r* = 0.95). Because total scan duration is the most commonly used heuristic for assessing PFM suitability, we next examined how expert judgments relate to scan duration. In contrast to NSI, total scan duration showed a more modest association with perceived dataset quality (**Fig. 5D**; r = 0.61) and substantial dispersion, particularly at shorter durations. Datasets with matched scan lengths (e.g., 15 or 45 minutes; dashed lines) spanned a wide range of quality ratings and usability endorsements, with both high-and low-rated datasets observed at the same durations. Together, these results indicate that scan duration alone is an incomplete proxy for PFM suitability, whereas NSI may provide a more consistent and interpretable summary of expert judgments of dataset quality.

To assess whether NSI captures explanatory information about expert judgments beyond intrinsic spatial properties of the data and the field’s prevailing heuristic for PFM suitability—scan duration—we fit a linear mixed-effects model pooling ratings from all six blinded raters. The model included NSI, spatial autocorrelation (Moran’s I), spectral slope, and total scan duration as fixed effects, with rater modeled as a random effect. Within this framework, NSI emerged as the strongest predictor of expert quality ratings (β = 2.60 rating units per SD increase in NSI, p < 0.001). Spectral slope showed a weaker but significant association (β = 0.63, p = 0.0038), whereas Moran’s I and scan duration did not explain additional variance after accounting for shared structure among predictors. Together, these results indicate that NSI subsumes information captured by generic spatial metrics while providing explanatory power beyond scan duration, supporting its interpretation as a compact and interpretable measure that aligns closely with expert judgments of PFM suitability.

### Predictive Utility of NSI for PFM Usability

Together, the analyses above establish that NSI closely aligns with expert perceptions of dataset quality on a continuous scale. We next examined whether this relationship has practical predictive utility—that is, whether NSI can prospectively identify whether a given dataset would likely be judged suitable for PFM by human experts. Using raters’ binary usability judgments, we therefore evaluated the ability of NSI to predict rater-specific usability decisions at the single-dataset level. ROC analysis showed strong and consistent discrimination across raters (**Fig. 5E**), with AUC values ranging from 0.81 to 0.98 (median = 0.93). At rater-specific optimal operating points defined by Youden’s *J*, NSI achieved high combined sensitivity and specificity (median *J* = 0.80), indicating effective separation of usable and non-usable datasets with low false-positive rates.

While ROC analysis shows that NSI separates usable from non-usable datasets, it does not tell users how confident they should be in the usability of any given dataset. To address this, we fit a calibrated logistic model relating NSI to the probability that a dataset would be judged usable for PFM (**Fig. 5F**). This model generalized to unseen raters and datasets under nested cross-validation, exhibiting low probabilistic error (median Brier score = 0.18) and high discrimination (median out-of-sample AUC = 0.92; **Supplementary Fig. 4**). The calibrated curve reveals a smooth transition from low to high usability probability across the intermediate NSI range where rater disagreement was greatest (**Fig. 5B**). Importantly, this formulation supports a probabilistic interpretation of NSI rather than a dichotomous threshold: for example, NSI values near 0.4 correspond to elevated uncertainty in expert consensus, whereas values above 0.5 are associated with high confidence of PFM usability across raters (e.g., ∼0.4 as marginal and ≥0.5 as generally favorable). Together, these analyses show that NSI can be translated into a calibrated, generalizable estimate of PFM usability that supports interpretable and principled decision-making.

### NSI Accounts for Differences in the Rate of FC Reliability Accumulation Across Individuals

The results in **Figures 3–5** establish NSI as an index of dataset quality for PFM. A distinct and practically important question is how NSI relates to the stability of individual-level FC estimates. Although within-individual FC reliability depends primarily on scan duration ^14,16,54^, the reliability achieved at a fixed scan duration can vary substantially across individuals and datasets. Prior work ^43,55^ has shown that acquisition approaches with improved BOLD contrast and denoising capabilities, such as multi-echo fMRI ^44^, exhibit better overall reliability, and in the present study these same datasets tend to exhibit higher NSI values (**Supplementary Figure 6**). We therefore hypothesized that differences in NSI would account for variability in FC reliability trajectories across individuals and datasets.

To test this hypothesis, we analyzed data from n = 31 highly sampled individuals, each with >240 minutes of motion-censored resting-state fMRI data (the same sample used in **Fig. 2**, separate from datasets used in **Fig. 5**). For each participant, progressively larger test subsets of motion-censored data (5–100 minutes) were selected, from which NSI was computed. To relate NSI to reliability, FC reliability was estimated from the same test subsets by comparing vertex-level FC matrices derived from each subset to a non-overlapping 120-minute reference epoch from the same individual. To reduce dependence on any particular session ordering, this procedure was repeated across multiple session-preserving circular rotations, such that different portions of each individual’s data alternately served as test and reference epochs. Because our hypothesis concerned whether NSI accounts for differences in the rate at which FC reliability accumulates across individuals, NSI was decomposed into a between-subject component (NSI_Between_; subject mean) and a within-subject component (NSI_Within_; duration-specific deviations).

To visualize the relationship between NSI and FC reliability prior to explicit modeling, we arranged subject-by-duration reliability estimates into a matrix ordered by NSI_Between_ (**Fig. 6A**). This representation revealed a clear gradient: at matched scan durations, individuals with higher NSI exhibited higher FC reliability, whereas low-NSI datasets showed slower gains and lower attainable reliability overall. To formally test this visually apparent pattern, we next fit linear mixed-effects models to the reliability-duration data. Reliability–duration curves differed significantly across NSI tertiles (**Fig. 6B**; linear mixed-effects model with subject as a random effect; main effect of NSI tertile: F(2,242) = 68.43, p < 0.001; NSI tertile × duration interaction: F(2,242) = 4.63, p = 0.01), indicating higher overall FC reliability and faster accumulation of reliable FC in high-NSI datasets.

**Figure 6.**
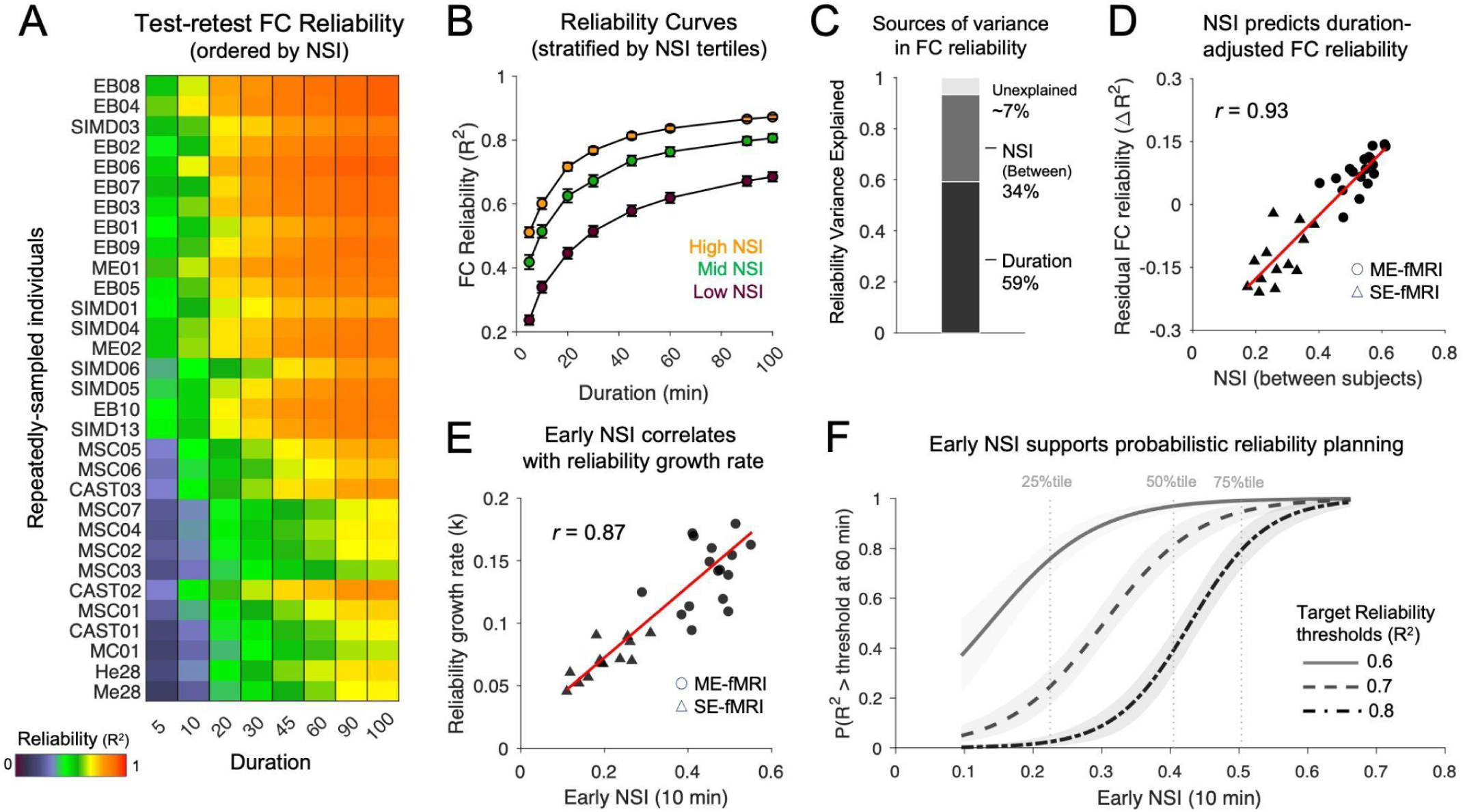
NSI accounts for individual differences in functional connectivity reliability trajectories. **(A)** Test–retest functional connectivity (FC) reliability (R²) across increasing scan durations (5–100 min) for repeatedly sampled individuals, ordered by Network Similarity Index (NSI). Warmer colors indicate higher reliability, revealing a clear gradient in achievable reliability as a function of NSI at matched durations. **(B)** Reliability–duration curves stratified by NSI tertiles (low, mid, high). Qualitatively, individuals with higher NSI exhibit more rapid accumulation of reliable FC estimates and higher asymptotic reliability, whereas low-NSI data show slower growth and lower ceilings. **(C)** Marginal pseudo-R² from mixed-effects models (fixed effects only, logit(R²) scale) summarizing the relative explanatory contributions of scan duration and NSI to FC reliability. Scan duration explains the majority of variance (59%), but between-subject differences in NSI (each subject’s mean NSI across durations) account for a substantial additional fraction of variance (34%) beyond duration alone. **(D)** Relationship between between-subject NSI (subject mean across durations) and duration-adjusted FC reliability (residuals from a duration-only model), demonstrating a strong positive association. Marker symbols denote acquisition sequence type (circles: multi-echo fMRI; triangles: single-echo fMRI), highlighting sequence-dependent differences that likely contribute to NSI-related reliability variation. **(E)** Early NSI (computed from the first 10 min of data) is correlated with individual differences in reliability growth rate (k), indicating that early data quality is associated with how quickly reliable FC estimates tend to accumulate with additional data. **(F)** Calibrated probabilistic models mapping early NSI (calculated using the first 10 minutes of data) to the probability of meeting or exceeding three reliability thresholds (R² = 0.6, 0.7, and 0.8) at 60 minutes of data. Vertical dashed lines denote the 25th, 50th, and 75th percentiles of early NSI observed in the empirical datasets, providing reference points for the range of early NSI values typically encountered in practice. Together, these curves illustrate how early data quality constrains the likelihood of achieving target FC reliability with additional scanning.

Having established this graded relationship using a categorical approximation of NSI, we next asked whether NSI and scan duration make independent and additive contributions to FC reliability. To address this question, we fit a series of linear mixed-effects models with reliability as the outcome, scan duration and NSI as fixed effects, and subject as a random effect to account for repeated measurements within individuals. A baseline model including scan duration alone showed a strong association between longer scans and higher FC reliability. Adding NSI_Between_ to this model significantly improved model fit (likelihood-ratio test: χ²(1) = 61.56, p < 0.001) and substantially increased the marginal explained variance from R² = 0.59 to R² = 0.93 (ΔR² = 0.34; **Fig. 6C**). In this combined model, both scan duration (β = 0.63, p < 0.001) and NSI_Between_ (β = 0.48, p < 0.001) were strong, independent predictors of FC reliability, indicating that between-subject differences in NSI account for substantial variability in reliability beyond scan time alone, even at matched durations.

To visualize the duration-adjusted contribution of NSI, we plotted subject-level gains in FC reliability relative to the duration-only model (ΔR²; observed minus duration-predicted reliability) against NSI_Between_, revealing a strong positive association (r = 0.93; **Fig. 6D**). Although this relationship was evident within both multi-echo (*r* = 0.67, p = 0.002) and single-echo (*r* = 0.59, p = 0.03) datasets when considered separately, multi-echo datasets tended to occupy higher ranges of both NSI_Between_ and duration-adjusted FC reliability, indicating that this difference in acquisition strategy contributes to but does not fully explain the NSI–reliability relationship.

### Reliability Growth and Decision Utility of NSI for FC Reliability

Having shown that higher NSI is associated with both higher overall FC reliability and faster accumulation of reliability with increasing scan duration, we next asked whether NSI measured from a single scan is informative about subsequent gains in reliability. We modeled how quickly FC estimates stabilize using a simple growth curve, in which a single parameter captures the speed at which reliability improves, independent of the maximum reliability eventually reached. Early NSI, computed from the first 10 minutes of data (a typical duration for a single fMRI run), was strongly correlated with this growth rate (*r* = 0.87; **Fig. 6E**), confirming that higher NSI is associated with greater scan efficiency—that is, faster gains in reliability per unit time—rather than differences in asymptotic reliability alone.

This relationship indicates that early NSI reflects not only contemporaneous data quality, but also how efficiently additional data contribute to improved reliability. **Figure 6F** illustrates an example decision-support use case in which NSI computed from a brief “scout” scan (10 minutes) is used to estimate the likelihood of achieving a target level of FC reliability at a given total scan duration. Across three reliability targets (R² = 0.6, 0.7, and 0.8), high early NSI values were associated with substantially greater likelihood of reaching the desired reliability threshold, whereas low-NSI datasets remained unlikely to do so even with additional scanning. Rather than providing point predictions for individual datasets, this framework translates early assessments of data quality into interpretable expectations about reliability accumulation, supporting principled decisions about data sufficiency and anticipated returns from further data collection across diverse acquisition contexts.

Leveraging this association, we developed a flexible set of calibrated probabilistic models that link NSI values to expected accumulation of FC reliability across scan durations. These models were evaluated under a stringent leave-one-subject-out framework with an additional leave-one-dataset-out constraint, ensuring complete separation of individuals and acquisition sites during training. To illustrate one representative use case, we report results for a model in which NSI from a 10-minute scout scan is used to estimate the likelihood of exceeding R² = 0.7 reliability at 60 minutes (**Fig. 6F**). This model showed low probabilistic error and strong discrimination, with median out-of-sample Brier scores across held-out sites ranging from 0.005 to 0.189 and a pooled out-of-sample AUC of 0.89 (see **Supplementary Fig. 5**). Importantly, this example reflects just one configuration within a broader modeling framework that supports prospective estimation across a range of scenarios, including projecting reliability at a given scan duration from its observed NSI, in addition to anticipating reliability gains with additional data collection. Together, these results demonstrate that NSI can be used to assess both the quality and reliability of FC, and to inform data collection decisions in precision fMRI studies.

## DISCUSSION

Investigators using PFM methods are often asked to define the data requirements for this approach. To date, experts have typically cited scan duration–based heuristics, recommending acquiring at least 30 or 40 minutes of data per subject ^14,16,43^, while often conceding that, in practice, the answer depends on factors like MRI acquisition protocol ^43,55^, or the degree of artifact burden and the effectiveness of denoising procedures ^43^. Implicit in these caveats is the recognition that scan duration or head motion estimates alone are incomplete proxies for whether a dataset expresses the large-scale functional organization required to support meaningful interpretation of individual-specific FC.

To address this challenge, we introduce NSI, a quantitative measure of the integrity of low–spatial-frequency network organization in individual fMRI datasets. Across multiple validation analyses, NSI covaried with independent spatial measures of large-scale coherence, decreased with targeted removal of BOLD-like signal, and closely tracked blinded expert judgments of PFM suitability. In repeatedly sampled individuals, NSI explained substantial between-subject differences in FC reliability at matched scan durations and indexed differences in the efficiency with which reliability accumulated as additional data were collected. Together, these findings position NSI as a concise and interpretable framework for evaluating individual-level FC data.

### A Shared Network Scaffold for Individual-Level Mapping

The central motivating observation of this work is that fine-scale individual-specific functional organization is expressed relative to a conserved, low–spatial-frequency network scaffold that is largely shared across individuals. This scaffold captures the dominant component of FC variance (**Fig. 2B**) and provides a stable reference frame within which finer-grained, person-specific features are reliably expressed. Conceptually, this organization is analogous to human facial anatomy. Across individuals, faces share a common large-scale structure, including a similar overall shape and a similar relative arrangement of core features, while individual identity emerges from finer-scale variations within that shared framework. In the brain, the shared low-frequency scaffold plays an analogous role, defining the broad outline of canonical large-scale network organization ^3,4^, while individual-specific features arise from reliable deviations in the size, shape, and boundaries of particular network representations ^56–58^.

This framework is conceptually related to prior work showing that functional brain networks are dominated by stable group- and individual-level factors rather than transient state or task effects ^32^. Whereas that work focused on parsing sources of FC variability across individuals, time, and cognitive states, here we focus on how functional organization is distributed across spatial scales within individual-specific FC patterns. By operating at the vertex level and explicitly decomposing spatial structure, we show that individual-level FC can be reliably separated into low–spatial-frequency and fine-scale components. Both components exhibit high split-half reliability within individuals (**Fig. 2C-D**). However, only the low–spatial-frequency component shows substantial similarity across individuals, whereas fine-scale structure is reliable within individuals yet largely idiosyncratic across individuals. Viewed in this way, the shared low–spatial-frequency component provides a common organizational reference for individual-specific FC patterns, rather than a constraint on individual variation itself. This distinction motivates the use of large-scale network structure, as quantified by NSI, as a basis for assessing the interpretability of individual-level functional maps while preserving meaningful fine-scale differences across individuals.

### Two Practical Applications of NSI for Precision fMRI

As a quantitative assay of the large-scale network organization required for interpretable individual-level FC, NSI affords two complementary, immediate applications for PFM studies: (i) retrospective or prospective screening of datasets for PFM suitability, and (ii) prospective planning of data collection based on the inferred reliability of the data already in hand and the expected reliability gains from additional data collection.

First, NSI provides a principled way to screen retrospective or ongoing datasets for PFM suitability. By calibrating NSI against blinded expert judgments, we show that NSI can be translated into a probabilistic screening tool that identifies datasets unlikely to support meaningful individualized network analyses (**Fig. 5**). This capability is particularly valuable for investigators new to PFM, for whom it can be difficult to determine whether a dataset expresses the coherent network structure required for individualized mapping. Rather than relying on scan duration alone, NSI offers a compact, interpretable summary of data quality that can guide transparent exclusion, stratification, or prioritization decisions before downstream modeling.

This potential screening role for NSI parallels practice in genomics, where measurements such as read depth and mapping quality are routinely used to determine whether sequencing data are sufficient for variant calling, with low-quality datasets filtered prior to inference ^59,60^. In clinical and translational settings, this screening role is particularly important when individualized connectivity maps are used to guide presurgical mapping ^61,62^ or personalized neuromodulation ^21,63^. In such contexts, low NSI values may provide an objective indication that patient-specific FC maps should be interpreted with caution, supporting conservative decisions to defer or down-weight individualized network information in favor of alternative approaches. Importantly, the level of evidence required for PFM usability is not uniform across applications. The amount of data quality and reliability needed depends on the scientific question and the network features of interest, with finer-grained or more individually variable properties demanding stronger evidence of signal integrity than broader, more conserved features. As a result, the same dataset may be sufficient for some individualized analyses but inadequate for others.

Second, NSI supports prospective planning by providing interpretable information about FC reliability at the scan duration from which it is computed, as well as expectations about how reliability is likely to accrue with additional data collection. Using NSI computed from a brief “scout” fMRI scan (e.g., 10 minutes), we show that investigators can indirectly estimate the likelihood of achieving a desired level of FC reliability at a given total scan duration (**Fig. 6**). This framework is particularly relevant for protocol development, study design, and grant planning, where decisions about scan duration and acquisition parameters must balance feasibility against reliability. Although prior work has emphasized total scan duration as a key design parameter alongside sample size ^64^, our results demonstrate that data quantity alone is an incomplete proxy for PFM suitability and test–retest reliability: even at matched durations, datasets can exhibit markedly different NSI values, corresponding to divergent expert judgments of PFM viability (**Fig. 5**) and distinct reliability trajectories (**Fig. 6**). In this context, NSI does not serve as an explicit measure of reliability itself, but rather as a proxy for underlying data quality that is systematically associated with how reliably FC estimates stabilize with additional data. In principle, FC reliability can be estimated more directly through formal statistical modeling of repeated measurements, though such approaches require assumptions and derivations that are distinct from the quality-based framework introduced here.

### NSI and Scan Duration Provide Complementary Information

Although validity and reliability are often treated as distinct measurement properties, signals that accurately reflect latent structure—such as well-constructed psychometric scales ^65^ or high-fidelity physiological recordings ^66^—typically reproduce more consistently. In the context of FC, the clear expression of canonical large-scale network organization provides an analogue of construct validity, reflecting the extent to which a dataset resolves known, functional networks rather than noise or artifact. Consistent with this principle, we found that NSI—an index of how clearly this canonical network structure is expressed in an individual fMRI dataset—explains substantial cross-subject variation in FC test–retest reliability at matched scan durations.

This relationship clarifies the distinct roles that scan duration and NSI play in shaping FC reproducibility. Scan duration primarily governs within-dataset gains in reliability as additional data accrue, but it does not explain why some datasets yield more reliable FC estimates than others despite extensive sampling. In contrast, NSI captures stable, subject-specific differences in achievable reliability at matched scan durations. Importantly, these differences persist among individuals scanned with identical acquisition and preprocessing, indicating that NSI indexes subject-specific data quality beyond known technical factors.

Viewed practically, these results motivate treating NSI as complementary to scan duration rather than as a replacement. Scan duration determines how reliability accumulates over time within a dataset, whereas NSI sets the effective gain of that accumulation across datasets. This distinction has direct implications for study design and data reuse, as NSI can guide both prospective selection of acquisition and preprocessing strategies likely to yield reliable inferences and retrospective prioritization of existing datasets suitable for individualized mapping.

### Signal Recoverability as a Complement to Artifact-Based QC

Quality control (QC) is central to fMRI analysis, with a longstanding emphasis on identifying, quantifying, and mitigating sources of artifact, especially head motion ^42^. Established measurements such as framewise displacement (FD) or DVARS ^67^, along with visualization strategies like grayplots ^68^, provide useful ways to characterize motion and its acute effects on the fMRI time series, as well as the global influences of other physiological fluctuations ^69,70^. However, these artifact-focused measures primarily index the potential burden of multiple discrete noise sources; they do not directly indicate whether meaningful network structure is actually discernible in an individual’s data after denoising.

NSI provides a complementary, signal-focused perspective. Consistent with this distinction, mean FD and other related head motion measurements explain only < 20% of variance in NSI values (**Fig. 3G**), and individuals with minimal motion nonetheless show a wide range of NSI, from low to high network identifiability. This indicates that data quality varies due to individual-specific factors not captured by head motion metrics alone. While NSI does not identify the specific sources of these quality differences, it is sensitive to their downstream consequences. By indexing the recoverability of BOLD-like, network-structured signal after denoising, NSI effectively summarizes how acquisition characteristics and preprocessing fidelity jointly shape the balance between structured signal and residual noise in an individual dataset.

### Practical Considerations for Interpreting NSI

First, as with any automated procedure, NSI should be viewed as an aid and not a replacement for careful manual inspection and expert judgment. NSI quantifies whether FC maps exhibit coherent, network-like spatial organization, but it is not designed to detect upstream failures that can invalidate FC estimation, such as errors in functional-to-anatomical co-registration or cortical surface reconstruction. This limitation was evident during the blinded rating process, in which an expert identified a pronounced surface registration failure in one dataset (“sub-AYHJR”) that did not materially affect NSI. Accordingly, NSI is intended to provide an objective and standardized assessment of network-level FC quality once preprocessing integrity has been established, replacing subjective judgments of FC rather than serving as a detector of all upstream processing failures.

Second, the approach presented here was developed and validated for surface-registered fMRI data represented in CIFTI format, with cortical hemispheres mapped to a standard surface mesh (fs_LR 32k) and combined with subcortical and cerebellar gray matter. NSI calculations require only that FC data and network priors be defined on a common spatial representation, and conversion to alternative surface meshes (e.g., fsaverage6) is straightforward. Given that the network FC priors emphasize low–spatial-frequency organization, the framework should also generalize naturally to volumetric representations, despite inter-individual variability in cortical folding ^71^. As such, the current surface-based implementation reflects a practical choice rather than a conceptual limitation of NSI, and analogous measures might, in principle, be defined for volumetric data.

## Conclusions

As interest in PFM continues to grow, clear guidance on when fMRI data are suitable for individualized mapping has lagged behind, with practice often relying on scan-duration heuristics or artifact-focused quality control. Here, we introduce NSI, an interpretable measure of how strongly an individual dataset expresses the coherent, large-scale network structures required for PFM. In this way, NSI assesses suitability for PFM by directly interrogating the functional properties of the dataset itself, providing information beyond indirect proxies based on scan duration or head motion, and thereby supporting more informed study design, protocol optimization, and transparent screening of legacy datasets for individualized mapping. Here, we provide an open-source codebase for NSI-based quality evaluation and precision fMRI analysis, along with expert-anchored usability ranges that offer a practical starting point for more standardized and reproducible application of PFM across studies and laboratories.

## Declaration of Generative AI and AI-assisted Technologies in the writing process

During the preparation of this work, the authors used ChatGPT (OpenAI) to assist with editing and organizing manuscript text for clarity and readability, and to aid in the organization and documentation of analysis code (e.g., commenting, refactoring for clarity and efficiency, and reviewing code for potential errors). All scientific content, analyses, interpretations, and conclusions were developed by the authors. The authors reviewed and edited all AI-assisted material and take full responsibility for the content of the published article.

## STAR Methods

### KEY RESOURCES TABLE

| REAGENT or RESOURCE | SOURCE | IDENTIFIER |
| --- | --- | --- |
| Deposited Data |  |  |
| PFM-Blind30 | This paper | https://openneuro.org/datasets/DS-XXXXXX |
| Software and Algorithms |  |  |
| PFM-NSI | This paper | https://github.com/cjl2007/XXXXXX |
| MATLAB | MathWorks | https://www.mathworks.com/ |
| Connectome Workbench | <sup>50</sup> | https://www.humanconnectome.org/software/connectome-workbench |
| Tedana | <sup>47</sup> | https://tedana.readthedocs.io/en/latest/ |
| FreeSurfer | <sup>72</sup> | https://surfer.nmr.mgh.harvard.edu/ |

#### Lead Contact

Further information and requests for resources should be directed to and will be fulfilled by the Lead Contact, Charles J. Lynch.

#### Materials Availability

This study did not generate new unique reagents. MATLAB code implementing all NSI-related procedures is being finalized for public release and will be made openly available via a GitHub repository upon publication, together with documentation describing software dependencies and execution order. During the peer-review process, access to the codebase is restricted to reviewers. Researchers interested in testing a pre-release (beta) version of the software during this period are welcome to contact the lead author to arrange access. De-identified CIFTI datasets used in the blinded rating experiment (PFM-Blind30) will be released on OpenNeuro upon publication. These data are withheld in the current preprint to preserve the option of recruiting additional blinded raters prior to final publication.

#### Overview of Experimental Procedures

Experimental procedures were designed to formalize manual inspection–based judgments of fMRI data suitability for PFM (**Fig. 1**), evaluate the assumptions underlying those judgments, and develop a quantitative measure of whether a dataset expresses the large-scale network organization required for interpretable individual-level inference. The work proceeded in four stages. First, we examined how functional connectivity (FC) structure is distributed across spatial scales and how individual-specific organization depends on the presence of shared large-scale network structure, using resting-state fMRI data from highly sampled individuals (**Fig. 2**). Second, motivated by this framework, we developed the Network Similarity Index (NSI) as an algorithmic analog of manual quality control, quantifying the extent to which seed-based FC maps express canonical large-scale functional network structure (**Fig. 3**). Third, we evaluated the construct validity of NSI across heterogeneous fMRI datasets by testing convergence with independent spatial metrics (Moran’s I and spectral slope), sensitivity to controlled denoising perturbations using multi-echo ICA (**Fig. 4**), and correspondence with blinded expert judgments of PFM dataset usability (**Fig. 5**). Finally, we assessed practical utility by jointly modeling scan duration and NSI in repeatedly sampled individuals to determine whether NSI accounts for between-subject differences in FC reliability beyond scan time alone and whether NSI indexes differences in the efficiency with which reliable FC estimates accumulate as additional data are acquired (**Fig. 6**).

### EXPERIMENTAL MODEL AND SUBJECT DETAILS

#### Participants

Participants were drawn from multiple independent neuroimaging datasets comprising both healthy and clinical populations. Across datasets, participants included healthy adults and adults with mood disorders (primarily major depressive disorder), as well as children and adolescents, including healthy youth from the ABCD study and children with autism spectrum disorder. This heterogeneity in demographic and clinical characteristics, paralleling the heterogeneity in acquisition protocols described in the following sections, is a deliberate design feature that allows us to evaluate NSI under conditions that approximate real-world research and clinical datasets. A summary of participant demographics for each dataset is provided in the **Supplementary Materials**.

### METHOD DETAILS

#### MRI image acquisition

MRI data were drawn from multiple independent datasets spanning healthy and clinical populations, with variation in scanner vendor (Siemens, GE, Phillips), field strength (3T and 7T), sequence type (single- and multi-echo EPI), and total scan duration. They range from modern, densely sampled longitudinal studies that have formed the backbone of current PFM literature (for example, the MyConnectome ^12,14^ and Midnight Scan Club ^16^) to legacy datasets with relatively short scan durations and older acquisition strategies. This heterogeneity is a design feature of the present study rather than a nuisance to be minimized. A detailed overview of MRI acquisition characteristics across all datasets, along with dataset-specific acquisition parameters and participant demographics, is provided in **Supplementary Fig. 6**.

#### Overview of MRI dataset processing

The datasets included in this study differ in acquisition provenance and preprocessing history. For approximately half of the datasets analyzed here, fMRI data were both acquired and processed using a common acquisition and preprocessing framework, and the procedures described in the following sections apply directly to these datasets (WCM-ME, WCM-LOD, WCM-ASD, WCM-TMS, SIMD). For a subset of additional datasets, raw data were acquired elsewhere but preprocessing was performed by the authors using closely related workflows (EuskalIBUR, SU-TMS, PNI), with minor modifications to accommodate differences in data acquisition (NSD, ABCD, 3D-TMS, UCSB). Finally, several datasets were obtained fully preprocessed from public repositories and were analyzed here largely as distributed, without reprocessing from raw data (MSC, CAST, MyConnectome). For clarity and brevity, the Methods below describe the core processing framework used for the majority of datasets, which reflects the dominant analytical approach in this study. Dataset-specific deviations from this framework are documented in the **Supplementary Materials**.

#### Anatomical preprocessing and cortical surface generation

Anatomical data were preprocessed and cortical surfaces generated using the Human Connectome Project (HCP) PreFreeSurfer, FreeSurfer, and PostFreeSurfer pipelines (version 4.3).

#### Multi-echo fMRI preprocessing

Preprocessing of multi-echo data minimized spatial interpolation and volumetric smoothing while preserving the alignment of echoes. The single-band reference (SBR) images (one per echo) for each scan were averaged. The resultant average SBR images were aligned, averaged, co-registered to the ACPC aligned T1-weighted anatomical image, and simultaneously corrected for spatial distortions using FSL’s topup and epi_reg programs. Freesurfer’s bbregister algorithm was used to refine this co-registration. For each scan, echoes were combined at each timepoint and a unique 6 DOF registration (one per volume) to the average SBR image was estimated using FSL’s MCFLIRT tool, using a 4-stage (sinc) optimization. All of these steps (co-registration to the average SBR image, ACPC alignment, and correcting for spatial distortions) were concatenated using FSL’s convertwarp tool and applied as a single spline warp to individual volumes of each echo after correcting for slice time differences using FSL’s slicetimer program. The functional images underwent a brain extraction using the co-registered brain extracted T1-weighted anatomical image as a mask and corrected for signal intensity inhomogeneities using ANT’s N4BiasFieldCorrection tool. All denoising was performed on preprocessed, ACPC-aligned images.

#### Multi-echo fMRI denoising

Preprocessed multi-echo data were submitted to multi-echo ICA (ME-ICA^46^), which is designed to isolate spatially structured T2*- (neurobiological; “BOLD-like”) and S0-dependent (non-neurobiological; “not BOLD-like”) signals and implemented using the “tedana.py” workflow^47^. In short, the preprocessed, ACPC-aligned echoes were first combined according to the average rate of T2* decay at each voxel across all time points by fitting the monoexponential decay, S(t) = S0e -t / T2*. From these T2* values, an optimally-combined multi-echo (OC-ME) time-series was obtained by combining echoes using a weighted average (WTE = TE * e -TE/ T2*). The covariance structure of all voxel time-courses was used to identify major signals in the OC-ME time-series using principal component and independent component analysis.

Components were classified as either T2*-dependent (and retained) or S0-dependent (and discarded), primarily according to their decay properties across echoes. All component classifications were manually reviewed by author CJL and revised when necessary following the criteria described in^73^. Mean gray matter time-series regression was performed to remove spatially diffuse noise. Temporal masks were generated for censoring high motion time-points using a framewise displacement (FD) threshold of 0.3 mm and a backward difference of two TRs, for an effective sampling rate comparable to historical FD measurements (approximately 2 to 4 seconds). Prior to the FD calculation, head realignment parameters were filtered using a stopband Butterworth filter (0.2 - 0.35 Hz) to attenuate the influence of respiration^74^ on motion parameters.

#### Surface processing and CIFTI generation of fMRI data

The denoised fMRI time-series was mapped to the individual’s fsLR 32k midthickness surfaces with native cortical geometry preserved (using the “-ribbon-constrained” method), combined into the Connectivity Informatics Technology Initiative (CIFTI) format using Connectome Workbench command line utilities^53^. This yielded time courses representative of the entire cortical surface, subcortex (accumbens, amygdala, caudate, hippocampus, pallidum, putamen, thalamus, brainstem), and cerebellum, but excluding non-gray matter tissue. Spurious coupling between subcortical voxels and adjacent cortical tissue was mitigated by regressing the average time-series of cortical tissue < 20 mm in Euclidean space from a subcortical voxel. All NSI and spatial coherence metrics (Moran’s I, spectral slope) were computed from data with no overt spatial smoothing. In contrast, FC reliability analyses were performed on spatially smoothed data, with Gaussian kernels applied in geodesic space for surface data and Euclidean space for volumetric data (σ = 1.7 mm).

#### Decomposition and Similarity Analysis of Functional Connectivity Across Spatial Scales

To decompose functional connectivity (FC) into shared large-scale and individual-specific components (**Main Text Fig. 2**), we performed a surface-based spatial frequency decomposition of seed-based FC maps using Laplacian eigenmodes defined on the fs_LR 32k cortical surface. Vertex-wise adjacency relationships were defined using a precomputed cortical neighbor table, from which a symmetric normalized graph Laplacian was constructed. The lowest nonzero Laplacian eigenmodes, corresponding to smoothly varying spatial patterns on the cortical surface, were obtained via sparse eigendecomposition. For each seed, FC maps were mean-centered and variance-normalized across vertices and projected into the Laplacian eigenbasis. The low–spatial-frequency component was reconstructed by retaining the first 400 nonzero Laplacian eigenmodes, capturing coarse, spatially smooth FC structure. The residual component was defined using the remaining higher-frequency modes up to the first 500 eigenmodes, isolating finer-scale spatial structure orthogonal to the low-frequency scaffold. For each map, the fraction of FC variance explained by the low-frequency and residual components was quantified as the proportion of total variance captured by each component.

To compare shared and individual-specific structure across spatial frequency bands, we quantified within- and between-subject similarity separately for low-frequency and residual FC maps. For within-subject comparisons, split-half FC maps were correlated across halves. For between-subject comparisons, FC maps from different individuals were compared using a best-matching seed strategy: for each seed in one subject, the maximum correlation with any seed in the other subject was identified, and similarity was summarized as the median of these maxima across seeds. This strategy avoids enforcing one-to-one correspondence between seed locations across individuals, which may map to different functional networks, and instead assesses similarity based on the presence of shared network-level FC profiles across seeds.

To assess whether observed within- and between-subject FC similarities exceeded those expected from generic spatial smoothness, we constructed a spatial null model based on random spherical rotations of cortical FC maps. For each permutation, FC maps from one subject were rotated on the cortical surface using precomputed vertex-wise rotation indices, preserving global spatial autocorrelation while disrupting vertex-to-vertex anatomical correspondence. Rotations were applied identically across all seeds, and unmapped vertices corresponding to the medial wall were filled using an independent alternate rotation to avoid structured missingness. Seed-by-seed similarity was then recomputed between unrotated and rotated maps, yielding a null distribution of best-matching seed correlations against which observed similarities were evaluated.

#### The Network Similarity Index (NSI)

The Network Similarity Index (NSI) was developed as a quantitative measure of fMRI dataset suitability for precision functional mapping (PFM). NSI quantifies the extent to which functional connectivity (FC) patterns within an individual express coherent large-scale brain network organization. For each seed region, the seed’s whole-brain FC map was modeled as a weighted combination of canonical large-scale functional network templates using ridge regression. In this formulation, the FC map serves as the response vector and the network templates as predictors. Ridge regularization mitigates collinearity among templates and accommodates overlapping network structure by allowing distributed combinations of templates to explain the observed FC pattern. Model performance was quantified using the coefficient of determination (R²), which defines the NSI value for each seed. Model fits were evaluated across a range of ridge penalties (λ = 1–50). Because NSI values were stable across this range (**Supplementary Fig. 2**), a value of λ = 10 was used for all subsequent analyses.

To summarize dataset-level quality, a global NSI value was computed as the median R² across all sampled seeds. Medians were used rather than means to provide robustness to skewed or multimodal seed-wise NSI distributions. By design, NSI evaluates whether FC patterns exhibit recognizable large-scale network organization independent of seed location. Accordingly, NSI does not assess whether a given seed matches an expected network identity at that location. This distinguishes NSI from correspondence-based approaches such as the Network Correspondence Toolbox (NCT ^75^), which quantify spatial overlap between empirical maps and predefined network atlases to support network labeling and interpretation. In contrast, NSI is intended as a continuous, dataset-level quality measure that assesses whether large-scale network structure is present at all.

NSI is calculated using the custom MATLAB function pfm_qc.m, which computes per-seed quality measurements and dataset-level summary scores, and pfm_qc_plots.m, visualizes seed-level distributions and, when supplied with the calibrated usability model (nsi_usability_model.mat), produces probabilistic estimates (with confidence intervals) of whether a dataset meets expert-defined criteria for PFM usability. In addition, the function conditional_reliability_from_nsi.m implements the NSI-based reliability model (nsi_reliability_model.mat), enabling estimation of expected FC reliability at a given scan duration and prospective evaluation of how reliability is likely to evolve with additional data collection based on NSI measured from an initial scan.

#### Sparse Whole-Brain Sampling

NSI were computed on a spatially sparse set of cortical vertices and subcortical voxels to maintain computational efficiency while preserving whole-brain coverage. For the cortex, adjacency relationships were defined using a table of vertex neighbors in the fs_LR 32k mesh, and a greedy selection algorithm iteratively added vertices only if none of their one-ring neighbors had already been selected. This yielded a uniformly distributed subset of cortical vertices with minimum one-ring spacing. For the subcortex, pairwise Euclidean distances between voxel coordinates were computed, and voxels within 2 mm of an already-selected voxel were excluded to maintain similar spatial sparsity. The resulting parcellation achieves near-uniform coverage of the brain while reducing the total number of seed locations by approximately fourfold, providing an efficient sampling framework for subsequent spatial measure computations.

#### Modeling NSI Associations with Spatial Measurements

To quantify the relationship between the Network Similarity Index (NSI) and independent measures of spatial organization (**Main Text Fig. 3**), we examined its association with Moran’s I (spatial autocorrelation) and spectral slope (spatial frequency content) using linear mixed-effects models implemented in MATLAB (fitlme, REML). For each participant, NSI, Moran’s I, and spectral slope were computed for every seed across the cortex. Moran’s I was computed by comparing FC values at each cortical vertex to those of its immediate neighbors on the surface mesh, aggregating these neighborwise similarities and normalizing by the overall variance of the FC map and the total adjacency weight. Spectral slope was computed by projecting each FC map onto cortical Laplacian eigenmodes, forming a spatial power spectrum, and estimating the slope of a linear fit between log power and log spatial frequency over an intermediate frequency range. To dissociate within- and between-subject sources of covariance, each spatial measurement was decomposed into a within-subject component (each seed’s deviation from the subject-specific mean) and a between-subject component (the subject-specific mean across all seeds). Both components were z-scored prior to modeling.

NSI was modeled as the dependent variable, with within- and between-subject components of Moran’s I or spectral slope entered as fixed effects and subject included as a random intercept to account for repeated seed-level observations within individuals. In Wilkinson notation, the primary models were: NSI ∼ 1 + X_within + X_between + (1 | Subject), where X denotes either Moran’s I or spectral slope. Marginal R² (variance explained by fixed effects) and conditional R² (variance explained by fixed plus random effects) were computed to summarize model fit.

#### Modeling Effects of Denoising Fidelity on NSI

To quantify the sensitivity of NSI to denoising fidelity (**Main Text Fig. 4**), we constructed a graded denoising-fidelity series within each individual run. This analysis used 100 adults with major depression, each completing four 15-min multi-echo resting-state runs processed with identical preprocessing and ME-ICA denoising. For each run, we began with the manually curated ICA decomposition, which provides discrete component labels (accepted/rejected) and continuous indices of component “BOLD-likeness” and “non-BOLD-likeness” (κ and ρ, respectively), along with component-wise variance contributions. We generated partially denoised variants by progressively swapping an increasing fraction of accepted components with an equal number of rejected components, holding the number of accepted components constant at each perturbation level. This manipulation simultaneously removes BOLD-like signals that would ordinarily be retained and reintroduces artifactual variance that would ordinarily be removed.

For each perturbation level, we summarized perturbation severity using manifest-derived variance-explained (VE) quantities that quantify (i) the effective fraction of BOLD-like variance removed (“RemovedBOLD”) and (ii) the effective fraction of non-BOLD-like variance introduced (“AddedNoise”). Although perturbation levels were defined by swapping equal numbers of BOLD-like and non-BOLD-like components, individual components differ substantially in the amount of variance they explain. Expressing perturbation severity in terms of VE therefore provides a continuous, normalized measure of the effective signal removal and noise injection, bounded between 0 and 100%, and comparable across runs and individuals. For each run and perturbation level, we calculated NSI. Effects of perturbation level were summarized as within-subject changes relative to the fully denoised baseline and standardized paired-difference effect sizes (Cohen’s d).

To evaluate which aspect of perturbation best explains NSI degradation, we fit linear mixed-effects models predicting within-subject changes in each metric (Δmetric = perturbed − baseline) from RemovedBOLD and AddedNoise, with a subject-specific random intercept. To make coefficients directly comparable, RemovedBOLD and AddedNoise were z-scored prior to modeling. We statistically compared the standardized coefficients by testing the linear contrast β(zRem) − β(zAdd) = 0 (Wald coefficient test) and computed a 95% confidence interval for the difference.

#### Expert Blind Ratings of Dataset Quality

To empirically validate whether NSI-PFM reflects expert judgments of data quality, we conducted a blinded rating study (**Main Text Fig. 5**). Twenty-five de-identified fMRI datasets were selected to span the full observed range of NSI values, approximating a normal distribution from low to high quality. Each subject’s functional connectivity (FC) maps were anonymized by zero-padding to conceal scan duration and replacing identifiers with randomized alphanumeric codes. Six expert raters (study co-authors ML, TL, EG, ZL, JD, and DP) with PFM experience independently reviewed all datasets in randomized order using Connectome Workbench, inspecting seed-based FC maps without access to acquisition details or metadata. Raters were unaware of the proposed NSI measure, its formulation, and all planned analyses at the time of evaluation and were only introduced to the NSI framework after ratings were completed.

For each dataset, raters provided (i) a continuous quality rating on a 1–10 scale and (ii) a binary judgment indicating whether the dataset was suitable for PFM (“Yes, usable” / “No, not usable”), along with optional qualitative notes. To assess intra-rater reliability, a fixed subset comprising 20% of datasets was duplicated and interleaved within the rating set; the same duplicated datasets were used for all raters, but the order in which datasets (including duplicates) were presented was independently randomized for each rater. Duplicate pairs were used exclusively to estimate within-rater consistency. For all inferential analyses relating ratings to NSI and scan duration, duplicate datasets were removed by retaining a single instance per duplicated dataset, randomly selected using a fixed seed for reproducibility.

#### Modeling of Expert Ratings and Usability Judgments

Associations between expert ratings and quantitative dataset measurements were evaluated using linear mixed-effects models to account for repeated ratings of the same dataset by multiple raters. Models included fixed effects for NSI as well as additional measurements (Moran’s I, spectral slope, and log-transformed scan duration). Dataset identity was included as a random intercept to account for multiple ratings of the same item, and rater effects were included either as random intercepts. The unique contribution of NSI was assessed by comparing the full model to a nested model omitting NSI using likelihood-ratio tests. Models were fit using maximum likelihood estimation. Binary usability judgments were analyzed separately to assess the ability of NSI to predict expert decisions regarding PFM suitability. Receiver operating characteristic (ROC) analysis was used to quantify discrimination between usable and unusable judgments.

#### Probabilistic Calibration of NSI to Usability Decisions

To translate NSI into an interpretable probabilistic decision aid, a binomial logistic calibration model was fit to map NSI values to the probability that a dataset would be judged usable for PFM. To evaluate generalization beyond the specific raters and datasets used for model fitting, calibration performance was assessed using a nested cross-validation scheme with joint rater and dataset holdout.

Six raters were included in the analysis. In each outer split, two raters were exhaustively held out (all 15 possible rater pairs), such that none of their ratings contributed to model training. Within each rater holdout split, datasets were further partitioned using 5-fold cross-validation, yielding training and test dataset subsets that were independent of the held-out raters. For each fold, the NSI-based binomial logistic calibration model was trained exclusively on ratings from the remaining four raters and the training datasets. Model fitting used only training-rater counts per dataset, with NSI centered using training data only. Model evaluation was then performed only on held-out datasets and held-out raters, producing out-of-sample predictions for rater–dataset combinations that were never observed during training. This design ensured that neither rater-specific decision thresholds nor dataset-specific characteristics could leak into model fitting. Across all rater splits and dataset folds, model performance was evaluated exclusively on held-out data by assessing calibration of predicted usability probabilities against empirical frequencies of held-out usability judgments, discrimination between usable and unusable judgments (area under the ROC curve), and probabilistic accuracy (Brier score). Performance measurements were summarized across all folds to quantify the extent to which NSI-based usability predictions generalized to unseen raters and unseen datasets, rather than reflecting overfitting to idiosyncratic rater behavior.

#### Progressive Estimation of NSI and Functional Connectivity Reliability

To quantify how data quality and scan duration jointly influence the reliability of functional connectivity (FC) estimates (**Main Text Fig. 6**), we computed both the Network Similarity Index (NSI) and FC reliability from progressively larger subsets of motion-censored resting-state fMRI data within each individual. For each subject, resting-state data were concatenated across sessions and motion censored using a framewise displacement threshold of 0.3 mm.

To reduce dependence on any particular session ordering, we employed a session-preserving rotation procedure. Time points were grouped by acquisition session, and multiple circular rotations of session order were generated by shifting session indices, with the number of rotations limited by the number of available sessions. Within each rotation, prefix subsets of increasing duration were extracted by selecting the first *N* time points corresponding to target durations (5, 10, 20, 30, 45, 60, 90, and 100 minutes). At each duration, FC reliability was quantified by comparing a “test” FC matrix computed from the selected subset to an independent, fixed-duration (120 minute) reference FC matrix estimated from a non-overlapping portion of the same individual’s data, drawn from the end of the rotated sequence. NSI was computed from the same motion-censored test subsets used for FC reliability estimation, ensuring direct correspondence between quality and reliability measures.

FC matrices were constructed using pairwise correlations between the same sparsely sampled seed set used for NSI (spanning cortical, subcortical, and cerebellar structures) and a distributed set of cortical surface targets, with spatially local cortical connections excluded using subject-specific geodesic distance matrices. Reliability was defined as the squared Pearson correlation (R²) between vectorized test and reference FC matrices. This metric indexes how well an FC estimate derived from a given amount of data recapitulates stable, individual-specific FC structure expressed in an independent reference dataset, providing a measure of FC stability as a function of scan duration.

#### Modeling of Scan Duration and NSI contributions to FC reliability

To quantify the relative contributions of scan duration and data quality to FC reliability, we fit linear mixed-effects models to subject-by-duration reliability estimates. Analyses were performed on the same progressive FC reliability values described above, using one observation per subject and duration. Because FC reliability values (R²) are bounded between 0 and 1, R² values were modeled on the logit(R²) scale. Scan duration was log-transformed to capture diminishing returns with additional data and standardized prior to modeling. Given the predominance of between-subject variance in NSI, NSI was decomposed into between-and within-subject components using person-mean centering.

NSI_Between_ was defined as each subject’s mean NSI across durations, mapped back to duration-specific rows, whereas NSI_Within_ captured deviations from that subject’s mean at each duration. NSI_Between_ was standardized at the subject level, and NSI_Within_ was standardized across rows. We fit nested linear mixed-effects models with a random intercept for subjects to account for repeated measures. The baseline model included scan duration only. Subsequent models added NSI_Between_ and NSI_Within_ as fixed effects. Model comparisons were performed using likelihood ratio tests on maximum-likelihood–fit models. Fixed-effect estimates were obtained from models refit using restricted maximum likelihood. To summarize the variance in FC reliability attributable to scan duration and NSI, we computed marginal pseudo-R² values based on fixed-effects-only predictions on the logit(R²) scale. This decomposition is intended to provide an interpretable summary of relative effect sizes rather than a formal partitioning of variance components.

#### Reliability Growth Curve Modeling

To characterize individual differences in the rate at which FC reliability accumulates with increasing scan duration, we fit an asymptotic growth model to each subject’s reliability–duration curve. Specifically, FC reliability was modeled as R(t) = Rmax × (1 − exp(−k × t)), where R(t) denotes FC reliability (R²) at scan duration t, Rmax represents the subject-specific asymptotic reliability ceiling, and k indexes the rate at which reliability increases with additional data. Model parameters were estimated separately for each subject by minimizing squared error between observed and predicted reliability values across available scan durations using nonlinear optimization.

#### NSI-based Modeling of Reliability Accumulation and Probabilistic Decision Framework

In the descriptive analyses above (mixed-effects modeling and growth curve modeling), NSI and FC reliability values were averaged across session-rotation iterations for each subject and duration. This approach emphasizes stable, dataset-level relationships while minimizing dependence on any particular temporal ordering of the data. In contrast, the analyses below operate at the level of individual rotation iterations, leveraging iteration-to-iteration variability in NSI and reliability accumulation to support probabilistic characterization of reliability outcomes. To model how early data quality relates to subsequent reliability accumulation, reliability growth parameters were estimated using an asymptotic growth model fit separately for each subject and each stochastic rotation iteration. This procedure yielded paired iteration-level estimates of NSI (computed from an initial segment of data) and FC reliability (R²). While many configurations were explored, including different early scan durations, total scan durations, and reliability thresholds, results are illustrated using one representative configuration in which early NSI was computed from the first 10 minutes of data.

Model performance was evaluated using a strict leave-one-subject-out (LOSO) cross-validation framework with an additional leave-one-dataset-out (LODO) constraint. For each held-out subject, all data from that subject and from other subjects acquired at the same site were excluded from model training. The relationship between early NSI and the reliability growth-rate parameter (k) was learned from the remaining subjects using linear regression applied to paired iteration-level observations pooled across training subjects. For held-out subjects, iteration-specific early NSI values were mapped to corresponding growth-rate estimates, while asymptotic reliability was fixed to the median value estimated from the training set.

Reliability at a target decision time was then computed using the fitted growth model and compared with the held-out subject’s observed reliability interpolated at the same duration, yielding an iteration-level binary outcome indicating whether a given reliability threshold was achieved. Although the primary figures focus on a representative decision time of 60 minutes and reliability thresholds of R² ≥ 0.6, 0.7, and 0.8, the same modeling framework supports alternative configurations, including different decision times, early NSI durations, and reliability criteria. These paired iteration-level outcomes were used to fit logistic regression models relating early NSI to the probability of exceeding each reliability threshold, providing an interpretable probabilistic decision framework. Model-predicted probabilities were evaluated across the observed NSI range, and uncertainty was quantified using bootstrap resampling to generate 95% confidence intervals. Generalization of this probabilistic framework was further assessed using LOSO+LODO cross-validation, with performance quantified using Brier scores, calibration curves, and receiver operating characteristic (ROC) analysis.

## Supplemental Materials

[Video S1 Link](https://wcm.box.com/s/o7ljl5sxhdgz8bfg31y11kq74lmdnrnu)

**Supplementary Video S1 Crawling seed-based functional connectivity in a high-quality individual resting-state fMRI dataset.** This video illustrates seed-based functional connectivity (FC) maps in a high-quality, individual dataset (ME01 ^43^; 6 hours of multi-echo resting-state fMRI), generated using wb_surfer2 (https://github.com/vanandrew/wbsurfer2). As the seed location systematically traverses the cortical surface, nearly every seed produces FC maps characterized by a spatially smooth, low–spatial-frequency structure that resembles a canonical large-scale functional network, with contiguous regions of positive and negative connectivity and minimal fine-grained speckling. This movie exemplifies the qualitative hallmark of datasets suitable for precision functional mapping (PFM), network-like organization that is robust to seed placement and interpretable across the cortex, even in regions typically vulnerable to susceptibility-related signal loss. Color scale indicates FC strength (warm = positive; cool = negative).

[Video S2 Link](https://wcm.box.com/s/b83kugiugx173u8r7u0uqw1buheybljx)

**Supplementary Video S2 Crawling seed-based functional connectivity in a lower-quality individual resting-state fMRI dataset.** This video illustrates seed-based functional connectivity (FC) maps in a qualitatively lower-quality individual dataset, again generated using wb_surfer2 (https://github.com/vanandrew/wbsurfer2). As the seed location systematically traverses the cortical surface, many seeds produce FC maps characterized by low-magnitude correlations and spatially fragmented, high–spatial-frequency structure. The resulting FC patterns frequently lack resemblance to canonical large-scale functional networks, instead exhibiting fine-grained speckling with rapid sign reversals and intermingled positive and negative values across neighboring vertices. This movie exemplifies the qualitative hallmark of datasets that are poorly suited for PFM: weak network organization that varies substantially with seed placement and is difficult to interpret across the cortex. Color scale indicates FC strength (warm = positive; cool = negative).

**Supplementary Figure 1.**
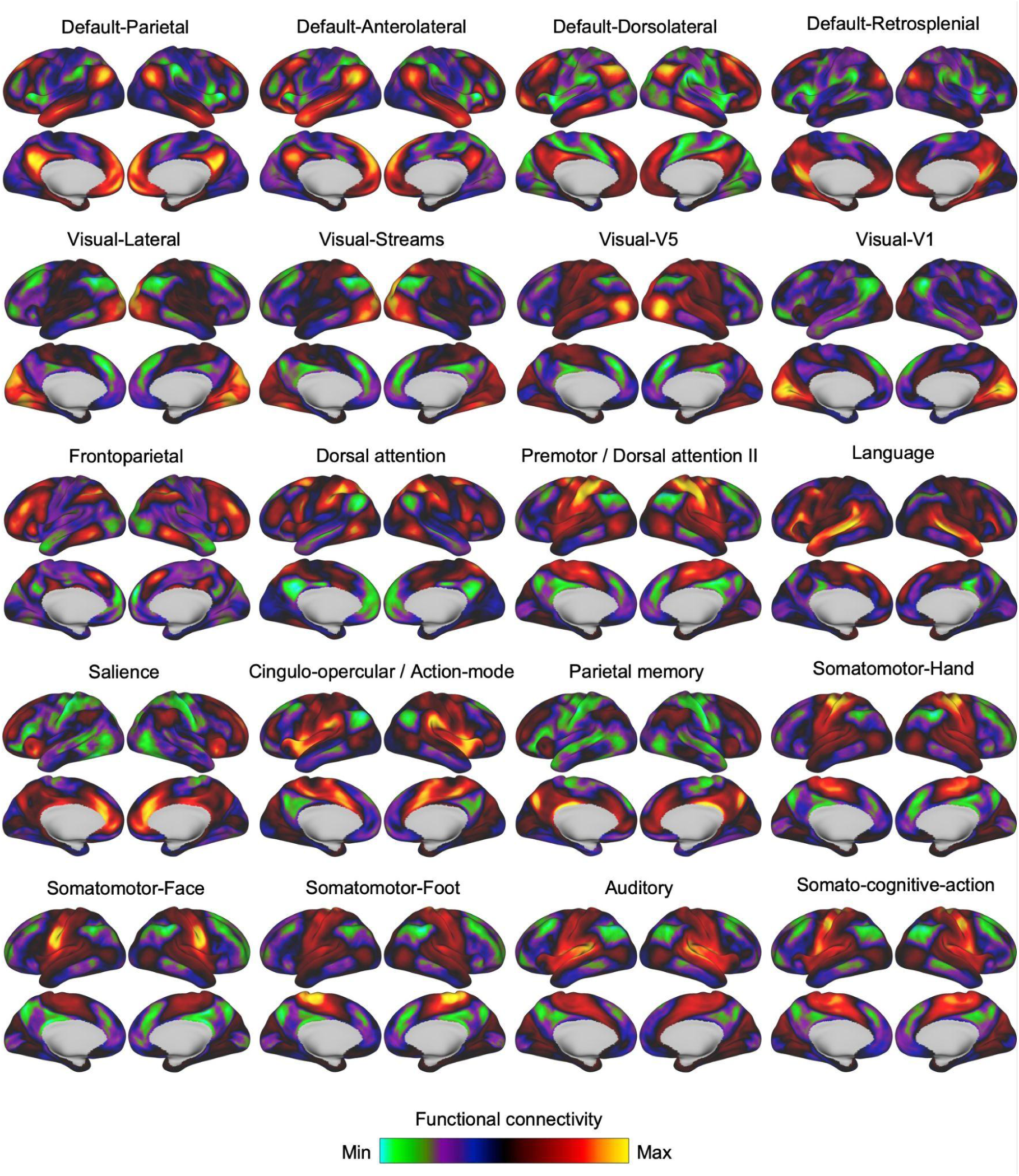
Default functional network priors (“WCM-20”) used for NSI calculations. Group-level functional network priors spanning canonical large-scale cortical systems were used as spatial references for NSI calculations. These priors were derived from a previous study ^17^, in which these functional networks were identified in N = 45 highly-sampled participants. For each individual, functional connectivity (FC) maps were computed for each identified network, and corresponding network maps were then averaged across individuals to generate group-level priors.The resulting WCM-20 priors include subdivisions of the default mode network, visual networks, frontoparietal and dorsal attention systems, language, salience and cingulo-opercular networks, parietal memory, somatomotor representations (hand, face, foot), auditory cortex, and somato-cognitive action networks.

**Supplementary Figure 2.**
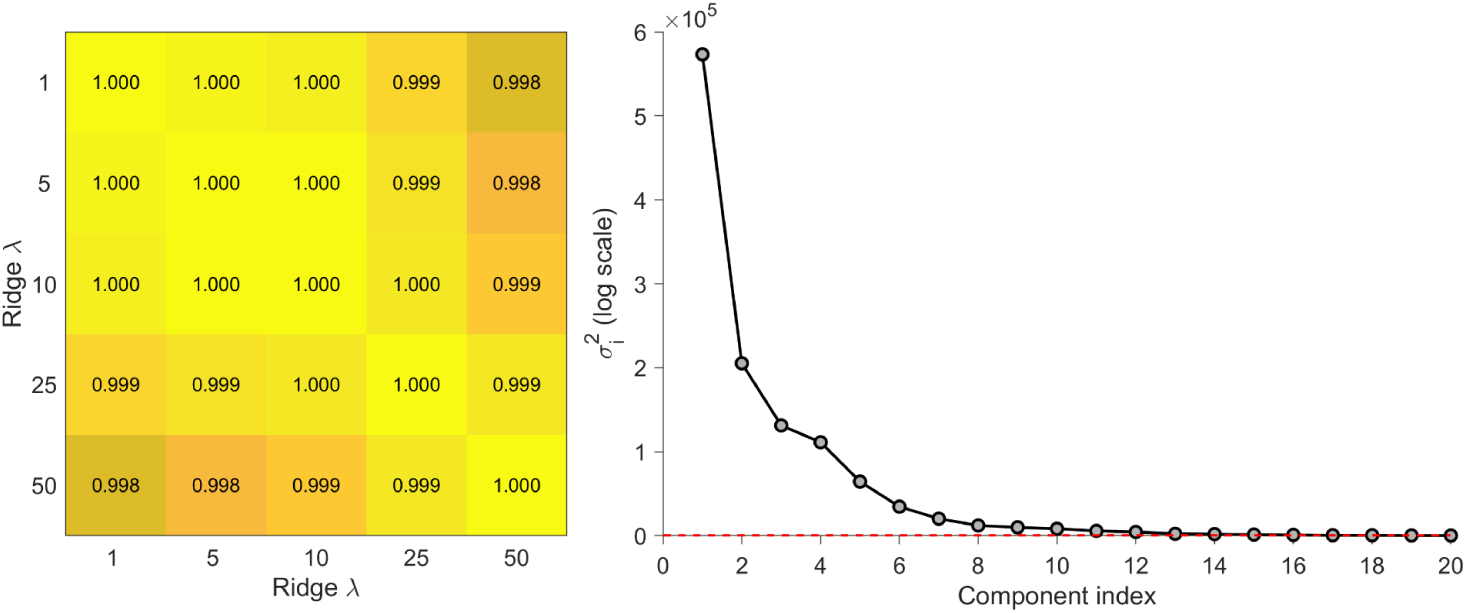
Robustness of the Network Similarity Index (NSI) to ridge regularization strength. (Left) Pairwise rank-order correlations (Spearman’s ρ) of subject-level median NSI values computed across a range of ridge regularization parameters (λ = 1–50). Rank ordering is nearly perfectly preserved across all λ values examined (ρ ≥ 0.998), indicating that relative dataset quality assessments are highly insensitive to the specific choice of regularization strength. (Right) Singular value spectrum of the functional network template matrix used in NSI computation, revealing a steep low-rank structure with rapidly decaying singular values. The λ values examined span a range comparable to—and extending beyond—the dominant singular values of the template matrix, covering regimes from weak to relatively strong regularization. In this context, ridge regularization primarily stabilizes inversion along low-variance dimensions, and performance degradation would be expected only for λ values substantially exceeding the leading singular values. Together, these results demonstrate that NSI estimates are robust across an appropriate and mechanistically motivated range of λ values, and that ridge regularization functions as a numerical stabilizer rather than a tuning parameter.

**Supplementary Figure 3.**
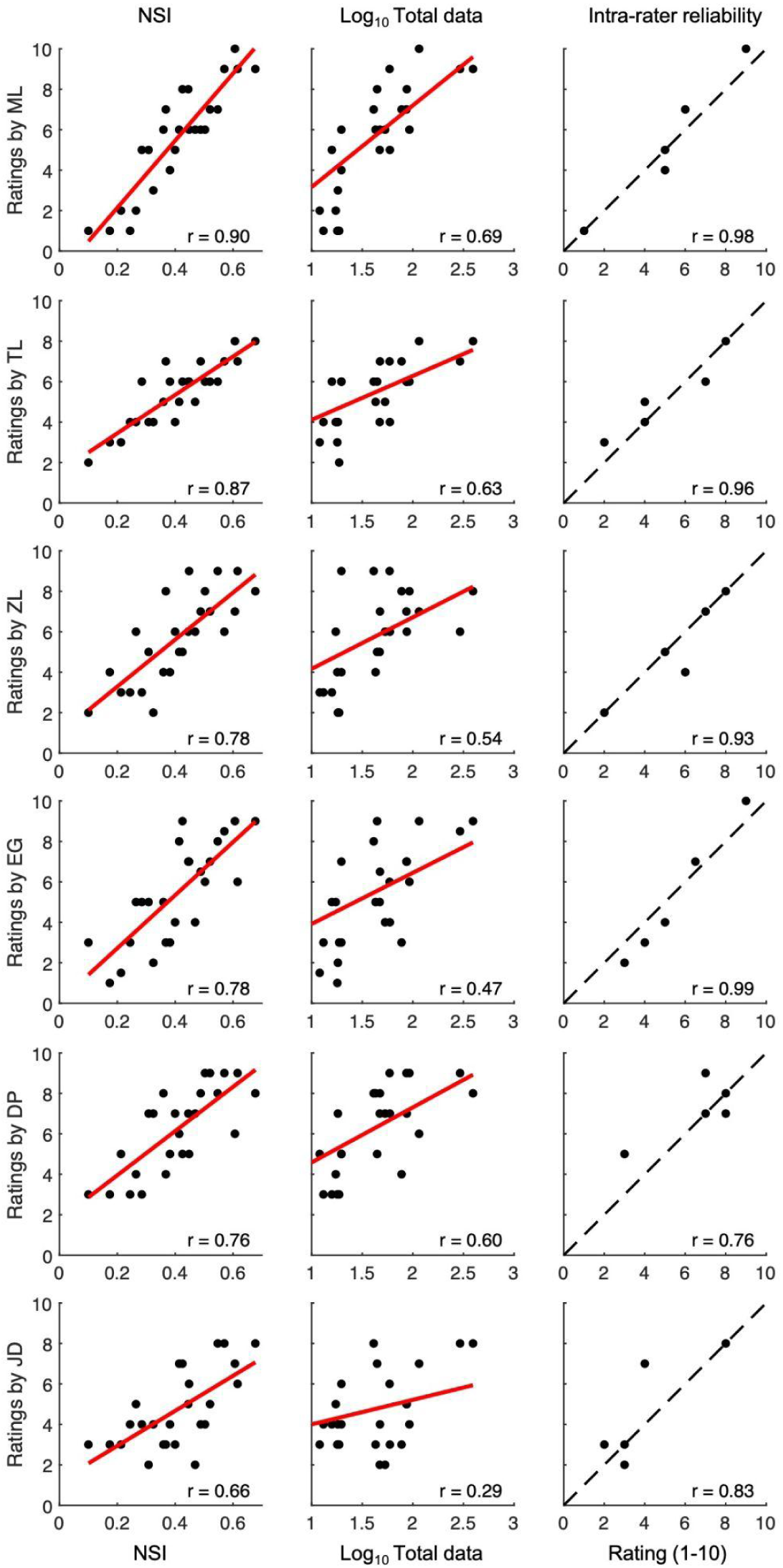
Relationships between NSI, total data per subject, and blinded expert judgments. Each row corresponds to a blinded human rater (n = 6), ordered by the strength of the association between rater scores and the Network Similarity Index (NSI). The first two columns show relationships between rater scores and NSI and total data duration (log-transformed). The third column shows intra-rater reliability estimated from duplicated datasets. In the intra-rater reliability column, paired ratings for duplicated datasets are plotted with the identity line (dashed black) shown for reference. Pearson correlation coefficients are reported in each panel. Across raters, NSI showed uniformly strong correspondence with expert judgments (r = 0.66–0.90, mean = 0.79), exceeding both the strength and consistency of associations with total data duration (r = 0.29–0.69, mean = 0.54). Intra-rater reliability was high overall (r = 0.76–0.99, mean = 0.91), indicating good consistency of expert ratings.

**Supplementary Figure 4.**
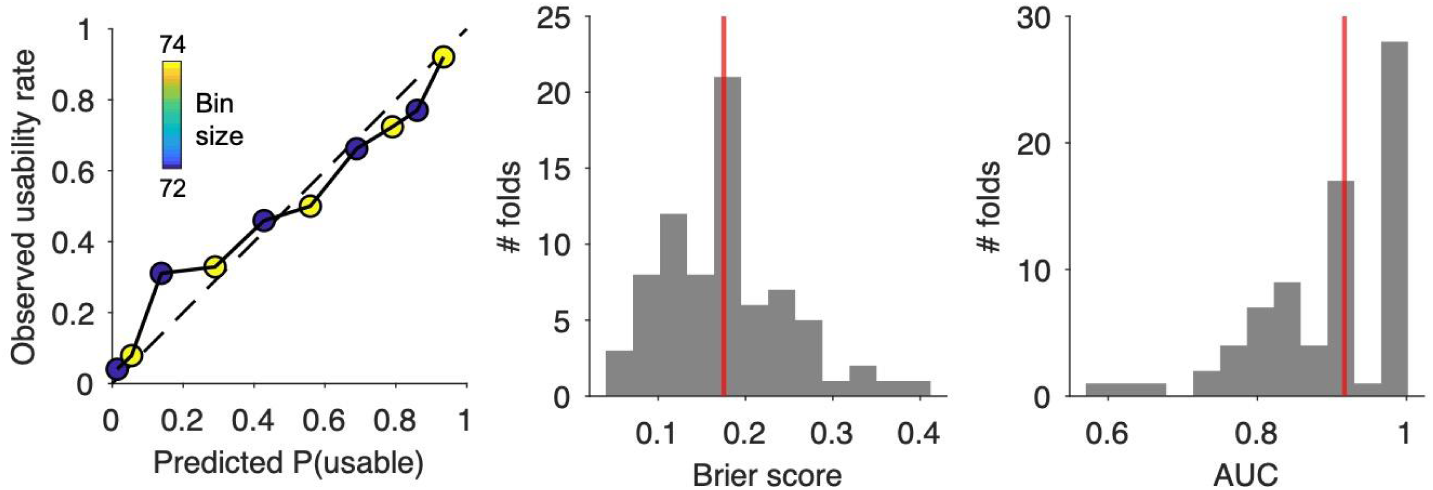
Cross-validation of NSI-based usability calibration. To assess whether the relationship between NSI and PFM usability generalizes beyond the specific raters and datasets used to estimate it, we performed a nested cross-validation procedure with joint rater and dataset holdout. Across all 15 possible rater splits, two raters were exhaustively held out in each outer split, and within each split datasets were further partitioned using 5-fold cross-validation. A binomial logistic calibration model (with a quadratic NSI term) was fit exclusively on ratings from the remaining raters and training datasets, and evaluated only on held-out datasets rated by held-out raters, ensuring strict separation between training and test data. Calibration of predicted PFM usability probabilities against observed held-out usability judgments, pooled across all held-out raters and datasets, shows close agreement with the identity line (Left), with points representing bins of out-of-sample predictions annotated by the number of observations per bin. Across cross-validation folds, probabilistic prediction error was low, as reflected by the distribution of Brier scores (median = 0.175; Middle), and discrimination between usable and unusable judgments was high, as reflected by the distribution of area under the ROC curve values (median AUC = 0.91; Right). Together, these results indicate that NSI-based usability predictions are well calibrated and generalize to unseen raters and datasets, rather than reflecting overfitting to idiosyncratic rater-specific decision thresholds.

**Supplementary Figure 5.**
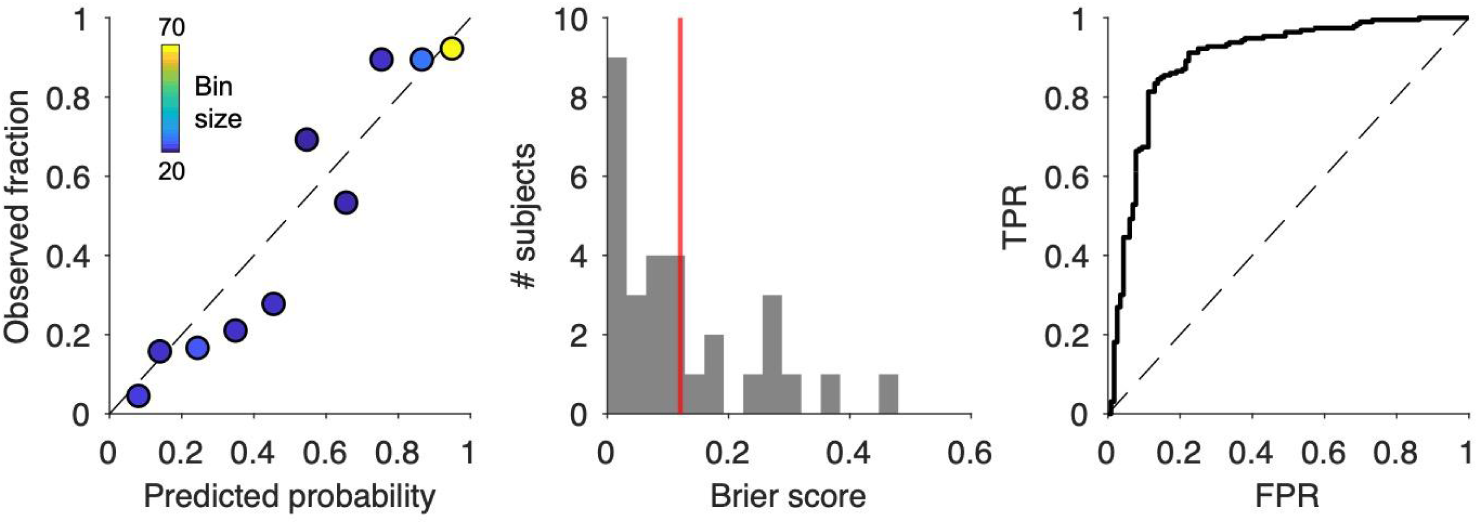
Cross-validation of NSI and FC reliability relationships. To assess whether early To evaluate whether the relationship between early Network Similarity Index (NSI) and subsequent functional connectivity (FC) reliability generalizes across individuals and acquisition contexts, we examined this relationship using a strict leave-one-subject-out (LOSO) cross-validation framework with an additional leave-one-dataset-out (LODO) constraint. Analyses shown here correspond to a representative decision-support configuration, in which NSI computed from an initial 10-minute “scout” scan is used to estimate the likelihood of exceeding a target FC reliability threshold (R² = 0.7) at a fixed decision time of 60 minutes.

In each outer split, all data from the held-out subject and all subjects acquired at the same site were excluded from model estimation. The association between early NSI (computed from the first 10 minutes of data) and the reliability growth-rate parameter (k) was estimated exclusively from the remaining subjects using paired iteration-level observations, while the asymptotic reliability parameter (Rmax) was fixed to the median value observed in the training set. This framework was used to assess how well early NSI supports probabilistic characterization of reliability outcomes at a fixed decision time (60 minutes), expressed as the likelihood of exceeding a target reliability threshold.

Calibration of out-of-sample predicted probabilities against observed reliability outcomes, pooled across all held-out subjects and datasets, shows close agreement with the identity line (Left), with points representing bins of predictions colored by bin size. Under strict LOSO+LODO evaluation, probabilistic prediction error remained low despite complete exclusion of entire datasets during training, with median Brier scores computed separately for each held-out site ranging from 0.005 to 0.189 (Weill Cornell Medicine, *n* = 8, median = 0.098; EuskalIBUR, *n* = 10, median = 0.009; Washington University, *n* = 10, median = 0.189; MyConnectome, *n* = 1, median = 0.013; UCSB, *n* = 2, median = 0.005; Middle; red line denotes the overall median). Discriminative performance summarized by the receiver operating characteristic curve pooled across held-out predictions (Right; dashed line indicates chance performance).

Importantly, across all cross-validation schemes, the probability of achieving the target reliability threshold increased monotonically with early NSI, indicating a stable and interpretable relationship between early network structure and future FC reliability. Together, these results demonstrate that NSI-based reliability estimates are reasonably calibrated and generalize across subjects and datasets, rather than reflecting overfitting to site- or cohort-specific characteristics.

**Supplementary Figure 6.**
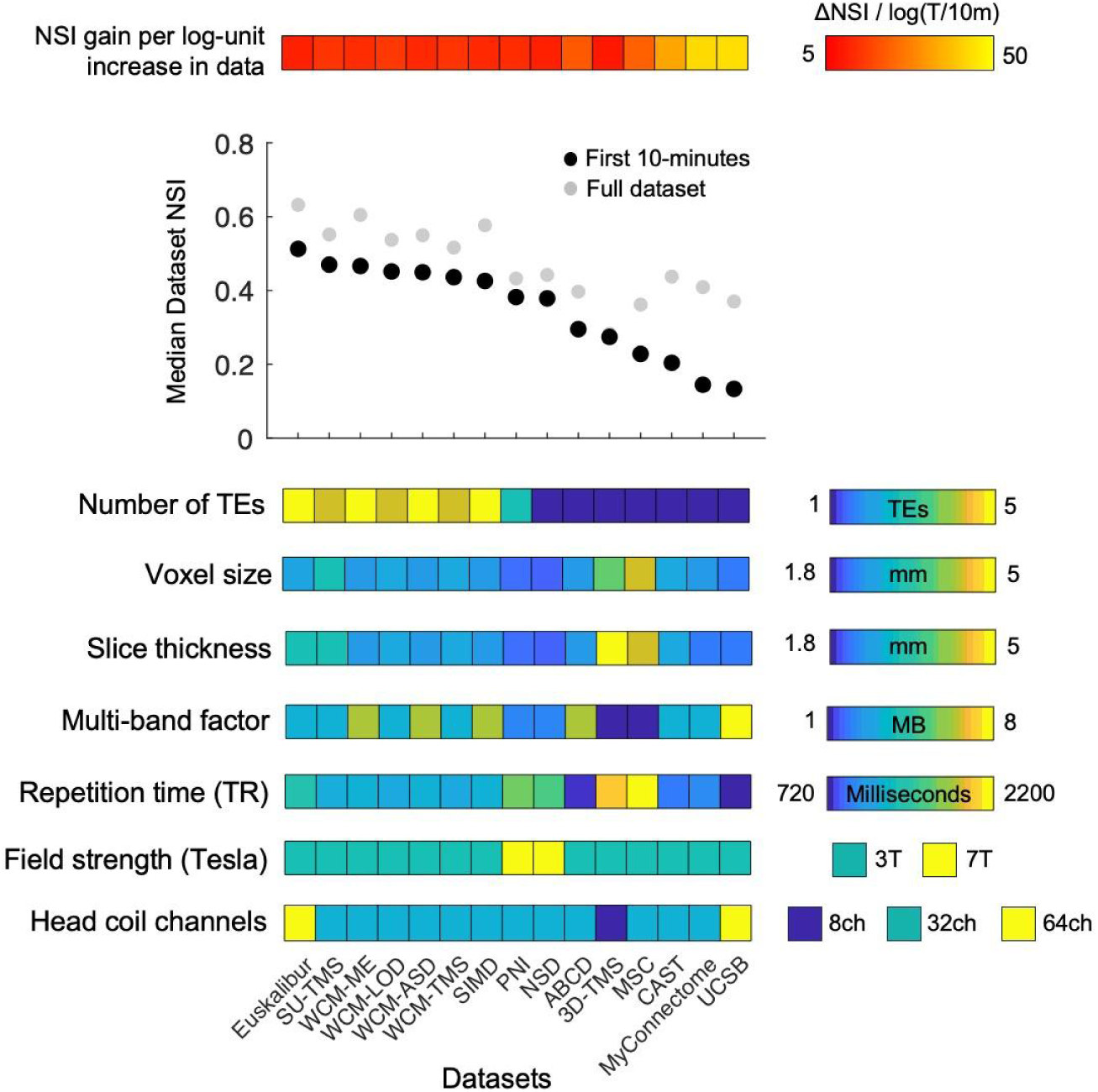
Network Similarity Index (NSI) across 15 different functional MRI datasets. Median NSI values computed across participants from the first 10 minutes of motion-censored, unsmoothed resting-state fMRI data are shown for each dataset (black markers). Gray markers indicate the corresponding dataset-level median NSI computed using all available data. By fixing scan duration for the 10-minute estimates, this analysis isolates differences attributable primarily to acquisition strategy and data processing, while comparison with full-dataset NSI illustrates how datasets differentially benefit from additional data. Datasets are ordered from left to right by their median 10-minute NSI. Selected acquisition parameters for each dataset are summarized below, including number of echo times (TEs), voxel size, slice thickness, multiband (MB) acceleration factor, repetition time (TR), field strength (3T or 7T), and head coil channel count (see next section for full acquisition and demographic details).

Across datasets, higher NSI values at fixed duration were most consistently observed in multi-echo acquisitions (TEs > 1), particularly those combining moderate spatial resolution (≈2.4–2.6 mm), moderate temporal resolution (TR ≈1300–1500 ms), and modest multiband acceleration (MB ≤ 4). In contrast, the lowest NSI values were observed in highly accelerated protocols (e.g., MB = 8, TR = 720 ms) with smaller voxel sizes (≈2 mm isotropic), consistent with the combined effects of aggressive acceleration and reduced voxel volume on BOLD contrast and signal-to-noise characteristics ^76^.

Comparison of 10-minute and full-dataset NSI values reveals substantial heterogeneity in the degree to which NSI increases with additional sampling (top heat strip). When normalized by the log-ratio of total scan duration to the initial 10-minute segment, datasets with lower initial NSI exhibited markedly larger NSI gains per log-fold increase in data than datasets with higher initial NSI (bottom three vs. top three datasets by 10-minute NSI; mean ΔNSI/log(T/10 min) ≈ 39.5 vs. 8.5). This pattern indicates that datasets with lower early NSI can go on to recover robust network-like structure with additional sampling, whereas datasets with higher early NSI exhibit more modest gains because they already have a well-resolved network organization.

### Supplemental Description of Functional MRI Datasets

The datasets used in this paper are described briefly below. We note that the Midnight Scan Club (MSC), MyConnectome, Cast Induced Plasticity, Natural Scenes Dataset (NSD), 28&Me, 28&He, ABCD, and Precision NeuroImaging and Connectomics (PNI) datasets were obtained online through various public repositories (e.g., https://openneuro.org/; https://aws.amazon.com/; https://osf.io/). Unless otherwise noted, all datasets were processed using the procedures described in the main text. Dataset-specific deviations from this processing framework are described where relevant.

#### EuskalIBUR (EB) dataset ^77^

Ten healthy adults (mean age = 31.4 ± 5.4 years; 5F/5M) were scanned at the Basque Center on Cognition, Brain and Language over ten weekly MRI sessions on a 3T Siemens PrismaFit scanner with a 64-channel head coil. In each session, T2*-weighted multi-echo BOLD fMRI was acquired with a multiband gradient-echo EPI sequence (TR = 1.5 s; TE₁–₅ = 10.60, 28.69, 46.78, 64.87, and 82.96 ms; flip angle = 70°; multiband factor = 4; GRAPPA = 2; 52 slices; voxel size = 2.4 × 2.4 × 3 mm³; FOV = 211 × 211 mm²; AP phase encoding). Each session comprised 64 minutes of multi-echo fMRI, including three task runs and four 10-minute resting-state runs; for the current work we used only the resting-state acquisitions, yielding approximately 407 ± 17.66 minutes of multi-echo resting-state fMRI data per participant across the study.

#### Stanford University TMS (SU-TMS) dataset ^17^

Adults undergoing TMS treatment were scanned at the Center for Neurobiological Imaging at Stanford University using a GE SIGNA 3T MRI system equipped with a Nova Medical 32-channel head coil. The sample comprised 42 participants (29 female, 13 male; mean age = 38.09 ± 12.77 years). Each participant completed two imaging visits, and at each visit four 7 min 30 s multi-echo, multi-band resting-state fMRI scans were acquired using a T2*-weighted echo-planar imaging sequence covering the full brain (TR = 1330 ms; TE₁–₄ = 13.7, 31.60, 49.50, and 67.40 ms; flip angle = 67°, corresponding to the Ernst angle for gray matter assuming T₁ = 1400 ms; field of view not reported; 3 mm isotropic voxels; 52 slices; anterior–posterior phase-encoding direction; in-plane acceleration factor = 2; multi-band acceleration factor = 4). This resulted in 60 minutes of resting-state fMRI data per participant (8 runs; 2,704 volumes total). Spin-echo EPI images with matched geometric parameters and opposite phase-encoding directions (AP and PA) were acquired prior to each resting-state scan for distortion correction. High-resolution T1-weighted and T2-weighted anatomical images were acquired at the end of each imaging session.

#### Weill Cornell Medicine Multi-echo (WCM-ME) dataset ^43^

Seven healthy adult men (mean age = 33.42 ± 9.10 years, 0F/7M) who underwent dense-sampling multi-echo resting-state fMRI at Weill Cornell Medicine. Data were acquired on a Siemens Magnetom Prisma 3T with a 32-channel head coil using the same multi-echo, multi-band T₂-weighted EPI sequence as the SIMD dataset (TR = 1355 ms; TE₁–₅ = 13.40, 31.11, 48.82, 66.53, 84.24 ms; flip angle = 68°; FOV = 216 mm; 2.4 mm isotropic voxels; 72 slices; in-plane acceleration factor = 2; multi-band factor = 6; AP phase encoding; 640 volumes per run, 14 min 27 s per scan). The number of resting-state runs varied substantially across participants and sessions, yielding 322.10 ± 312.08 minutes of multi-echo fMRI data per participant.

#### Weill Cornell Medicine Late-onset Depression (WCM-LOD) dataset ^17^

Five adults with late-onset major depressive disorder (mean age = 66.60 ± 5.31 years, 0M/5F) recruited at Weill Cornell Medicine, each contributing 59.05 ± 6.55 minutes of resting-state fMRI data. MRI data were acquired on a Siemens Magnetom Prisma 3T scanner at the Citigroup Biomedical Imaging Center using a Siemens 32-channel head coil. At each study visit, two multi-echo, multi-band resting-state fMRI scans were collected using a T2*-weighted EPI sequence covering the whole brain (TR = 1300 ms; TE₁–₄ = 12.60, 29.51, 46.42, 63.33 ms; FOV = 216 mm; flip angle = 67°; 2.5 mm isotropic voxels; 60 slices; AP phase-encoding; in-plane acceleration factor = 2; multi-band factor = 4; 480 volumes per scan; 10 min 38 s per run). Spin-echo EPI images with opposite phase-encoding directions (AP and PA) but identical geometry and echo spacing were acquired before each resting-state scan. Multi-echo T1-weighted (TR/TI = 2500/1000 ms; TE₁–₄ = 1.7, 3.6, 5.5, 7.4 ms; FOV = 256 mm; flip angle = 8°; 208 sagittal slices; 0.8 mm slice thickness) and T2-weighted anatomical images (TR = 3200 ms; TE = 563 ms; FOV = 256.

#### Weill Cornell Medicine Autism Spectrum Disorder (WCM-ASD) dataset ^17^

Six pediatric participants with autism spectrum disorder (ASD; ages 9–12 years), each contributing 2.94 ± 0.89 hours of multi-echo resting-state fMRI data. MRI scans were acquired on a Siemens Magnetom Prisma 3T scanner at the Citigroup Biomedical Imaging Center using a Siemens 32-channel head coil and the same multi-echo, multi-band T₂*-weighted EPI sequence as the WCM-ME adult sample (TR = 1355 ms; TE₁–₅ = 13.40, 31.11, 48.82, 66.53, 84.24 ms; flip angle = 68°; FOV = 216 mm; 2.4 mm isotropic voxels; 72 slices; in-plane acceleration factor = 2; multi-band factor = 6; 640 volumes per run; 14 min 27 s per scan). To minimize head motion, participants’ heads were stabilized using custom 3D-printed foam headcases throughout data collection.

#### Weill Cornell Medicine TMS (WCM-TMS) dataset ^17^

Adults with major depressive disorder drawn from two rTMS treatment studies at Weill Cornell Medicine (combined mean age = 42.62 ± 14.11 years; N = 48; 24M/24F; 58.86 minutes of multi-echo fMRI data per participant). MRI data were acquired on a Siemens Magnetom Prisma 3T scanner at the Citigroup Biomedical Imaging Center using a 32-channel head coil. At each visit, two multi-echo, multi-band resting-state fMRI scans were collected using a T2*-weighted EPI sequence (TR = 1300 ms; TE₁–₄ = 12.60, 29.51, 46.42, 63.33 ms; FOV = 216 mm; flip angle = 67°; 2.5 mm isotropic voxels; 60 slices; AP phase encoding; in-plane acceleration factor = 2; multi-band factor = 4; 650 volumes per scan, 14 min 5 s). Spin-echo EPI images with AP/PA phase-encoding were acquired for distortion correction, along with multi-echo T1-weighted and T2-weighted anatomical images using matching high-resolution parameters.

#### Serial Imaging of Major Depression (SIMD) ^17,21^

Eight adults with mood disorders completed repeated MRI sessions at the Citigroup Biomedical Imaging Center of Weill Cornell’s medical campus. The cohort included six individuals with unipolar major depression (mean age = 29.47 ± 8.28 years; 3F/3M) and two individuals with bipolar disorder (2M), all scanned using the same protocol. Data was acquired on a Siemens Magnetom Prisma 3T scanner with a Siemens 32-channel head coil. At each visit, two multi-echo, multi-band resting-state fMRI scans were collected using a T2*-weighted echo-planar sequence covering the full brain (TR = 1355 ms; TEs = 13.40/31.11/48.82/66.53/84.24 ms; FOV = 216 mm; flip angle = 68°; 2.4 mm isotropic voxels; 72 slices; AP phase-encoding; in-plane acceleration factor = 2; multiband factor = 6), with 640 volumes per scan (14 min 27 s each), yielding ∼29 minutes of resting-state data per visit. Across the six unipolar depression cases, participants contributed on average 621.49 ± 691.53 minutes of multi-echo fMRI data, and the two participants with bipolar disorder contributed 5.30 and 28.83 hours of multi-echo fMRI data per subject, respectively.

#### Precision NeuroImaging and Connectomics (PNI) dataset ^78^

Ten healthy adults (4 male, 6 female; mean age = 26.6 ± 4.6 years; 2 left-handed) with no history of neurological or psychiatric illness completed three MRI sessions at the McConnell Brain Imaging Centre (Montreal Neurological Institute). Imaging was performed on a Siemens MAGNETOM Terra 7T scanner using a 32-channel receive / 8-channel transmit head coil in parallel transmission mode. Each ∼90-min session combined high-resolution multimodal structural imaging (0.5–0.7 mm MP2RAGE T1 relaxometry, diffusion MRI with multi-shell acquisition, magnetization transfer, and multi-echo T2*-weighted imaging) with multi-echo BOLD fMRI acquired using a 2D multi-echo EPI sequence from CMRR (TR = 1690 ms; TEs = 10.8/27.3/43.8 ms; flip angle = 67°; 1.9 mm isotropic voxels; 75 slices; multiband factor = 3; FOV = 224 × 224 mm²). Across sessions, this protocol yielded a total of approximately 48 minutes of multi-echo fMRI data per subject.

#### Natural Scenes Dataset (NSD) ^79^

Eight healthy adults (mean age = 26.50 ± 4.24 years; 6F/2M) participated in ultra–high-field imaging at the University of Minnesota’s Center for Magnetic Resonance Research. Functional data were acquired on a 7T scanner using gradient-echo EPI with whole-brain coverage (including cerebellum; 84 axial slices; 1.8 mm isotropic voxels; FOV 216 × 216 mm²; matrix 120 × 120; TR = 1,600 ms; TE = 22.0 ms; flip angle = 62°; anterior–posterior phase-encoding; multiband acceleration factor = 3). Resting-state runs (188 volumes; ∼5 min each) were collected before and/or after the core NSD task runs in a subset of 7T sessions using the same acquisition parameters. Across the runs included in our analyses, this yielded a total of 115.17 ± 35.54 minutes of resting-state fMRI data per subject. Functional data were preprocessed by the original study authors ^79^, after which we performed denoising using ICA-AROMA ^80^ followed by mean gray matter time-series regression (MGTR). Because the functional data had already undergone a single interpolation step as part of standard preprocessing procedures (e.g., motion correction and distortion correction), and because the corresponding anatomical data had not been registered to functional space at that stage, we avoided additional resampling of the functional time-series by rigidly aligning FreeSurfer-derived cortical surfaces to the functional data. The denoised fMRI time-series was then mapped to each individual’s fsLR 32k midthickness surface using a ribbon-constrained procedure that preserved native cortical geometry and combined into Connectivity Informatics Technology Initiative (CIFTI) format using Connectome Workbench command-line utilities ^53^.

#### Adolescent Brain Cognitive Development (ABCD) dataset ^81^

We analyzed a subset of 100 children from the ABCD Study, drawn from our previous work (mean age = 9.46 ± 0.50 years; equal numbers of boys and girls). MRI data were collected across 21 sites on harmonized 3T systems (Siemens Prisma, GE 750, and Philips scanners) using multi-channel head coils. Functional images were acquired with a T2*-weighted multiband EPI sequence (TR = 800 ms; TE = 30 ms; flip angle = 52°; 2.4 mm isotropic voxels; 60 slices; multiband acceleration factor = 6). For the present work, we combined all available resting-state and task fMRI runs to increase the effective data per participant, yielding a total of 96.43 ± 15.02 minutes of fMRI data per-subject. Preprocessing followed the procedures described in the main text, adapted for single-echo acquisitions. Following preprocessing, data were denoised using ICA-AROMA ^80^ and MGTR. The denoised fMRI time-series was mapped to the individual’s fsLR 32k midthickness surfaces with native cortical geometry preserved (using the “-ribbon-constrained” method), combined into the CIFTI format using Connectome Workbench command line utilities^53^.

#### Three-D (3D-TMS) dataset ^82^

Patients with treatment-resistant major depressive disorder from the THREE-D randomized clinical trial who completed a 10-minute resting-state fMRI scan at baseline. Imaging was performed on a 3T GE HDx MRI system with an 8-channel phased-array head coil. For each participant, we acquired a high-resolution T1-weighted structural scan (fast spoiled gradient echo; TE = 12 ms, TI = 300 ms, flip angle = 20°, 116 sagittal slices, 1.5 mm thickness, no gap, 256 × 256 matrix, FOV = 240 mm) and a 10-minute T2*-weighted resting-state EPI scan (TE = 30 ms, TR = 2000 ms, flip angle = 85°, 32 axial slices, 3.5 x 3.5 x 5 mm voxels, 64 × 64 matrix, FOV = 220 mm). Preprocessing followed the procedures described in the main text, adapted for single-echo acquisitions. Preprocessed data were denoised using ICA-AROMA ^80^ and MGTR. The denoised fMRI time-series was mapped to the individual’s fsLR 32k midthickness surfaces with native cortical geometry preserved (using the “-ribbon-constrained” method), combined into the CIFTI format using Connectome Workbench command line utilities^53^.

#### Midnight Scan Club (MSC) dataset ^16^

Ten healthy adults (age = 29.1 ± 3.3 years; 5F/5M) were scanned at Washington University School of Medicine on a Siemens Trio 3T MRI scanner. Resting-state fMRI was collected across 10 late-night sessions on separate days, with each session including 30 minutes of eyes-open rest while participants fixated on a central crosshair. Resting-state images were acquired using a gradient-echo EPI sequence (TR = 2200 ms; TE = 27 ms; flip angle = 90°; 4 × 4 × 4 mm³ voxels; 36 slices). Across sessions, this protocol yielded a total of 300 minutes of resting-state fMRI data per participant. Because the surface-registered MSC data available on OpenNeuro are already spatially smoothed, denoised volumetric fMRI data were instead obtained and projected to the the individual’s fsLR 32k midthickness surfaces with native cortical geometry preserved (using the “-ribbon-constrained” method), combined into the CIFTI format using Connectome Workbench command line utilities^53^.

#### Cast-induced Plasticity dataset (CAST) ^38^

Three healthy right-handed adults (ages 25, 27, and 35 years; 2M/1F) underwent intensive longitudinal imaging at Washington University School of Medicine while their dominant upper extremity was immobilized in a cast for two weeks. Two participants were also scanned in the Midnight Scan Club (MSC02 and MSC06), allowing us to examine how changes in scanner hardware and pulse sequence affect data quality across studies. Participants were scanned daily for 42–64 consecutive days, with each session including a 30-min resting-state fMRI run (eyes open, fixation), yielding approximately 21–32 hours of rs-fMRI data per participant. BOLD data were acquired with gradient-echo EPI: one participant was scanned on a Siemens Trio 3T using single-band acquisition (TR = 2200 ms; TE = 27 ms; flip angle = 90°; 4 × 4 × 4 mm³ voxels; 64 × 64 × 36 matrix), while the later control sessions for this participant and all scans for the remaining two participants were collected on a Siemens Prisma 3T using higher-resolution multiband EPI (TR = 1100 ms; TE = 33 ms; flip angle = 84°; 2.6 mm isotropic voxels; 90 × 90 × 56 matrix; multiband factor = 4). Because the surface-registered CAST data available on OpenNeuro are already spatially smoothed, denoised volumetric fMRI data were instead obtained and projected to the the individual’s fsLR 32k midthickness surfaces with native cortical geometry preserved (using the “-ribbon-constrained” method), combined into the CIFTI format using Connectome Workbench command line utilities^53^.

#### MyConnectome (MC) dataset ^12^

A single healthy adult participant (45-year-old right-handed Caucasian male) was scanned over ∼18 months at the University of Texas at Austin using a Siemens Skyra 3T scanner with a 32-channel head coil. Resting-state fMRI was acquired in up to three sessions per week using a multi-band gradient-echo EPI sequence (TR = 1,160 ms; TE = 30 ms; flip angle = 63°; voxel size = 2.4 × 2.4 × 2.0 mm³; 68 slices; 96 × 96 matrix; FOV ≈ 230–240 mm; multiband factor = 4; 10-min scan length), with slices oriented ∼30° back from the AC–PC line. After excluding pilot sessions, 75 ten-minute resting-state runs were retained, yielding a total of 740 minutes of resting-state fMRI data from this single individual. Because surface-registered MyConnectome data on OpenNeuro are spatially smoothed and denoised volumetric data were not made available, raw fMRI data were obtained and preprocessed using the main-text pipeline adapted for single-echo acquisitions, with subsequent denoising using an aCompCor ^83^ framework. The denoised fMRI time-series was mapped to the individual’s fsLR 32k midthickness surfaces with native cortical geometry preserved (using the “-ribbon-constrained” method), combined into the CIFTI format using Connectome Workbench command line utilities^53^.

#### 28&ME dataset ^36^

A single right-handed woman (age 23) scanned daily across two 30-day studies (Study 1: natural menstrual cycle; Study 2: oral contraceptive use). Each session included 10 minutes of resting-state fMRI on a Siemens 3T Prisma with a 64-channel head coil using a T2*-weighted multi-band EPI sequence (72 oblique slices; TR = 720 ms; TE = 37 ms; voxel size = 2 mm³; flip angle = 56°; multiband factor = 8), preceded by high-resolution T1-weighted MPRAGE and fieldmap acquisition and accompanied by daily hormone assays and mood/lifestyle questionnaires. Head motion was minimized with a custom 3D-printed headcase for most sessions, yielding 60 sessions and approximately 600 minutes of resting-state fMRI data in total. Raw fMRI data were obtained and preprocessed using the main-text pipeline adapted for single-echo acquisitions, with subsequent denoising using an aCompCor ^83^ framework. The denoised fMRI time-series was mapped to the individual’s fsLR 32k midthickness surfaces with native cortical geometry preserved (using the “-ribbon-constrained” method), combined into the CIFTI format using Connectome Workbench command line utilities^53^.

#### 28&HE dataset ^37^

A single right-handed Caucasian male (age 26 years) without neuropsychiatric, endocrine, or head trauma history underwent intensive longitudinal imaging over 30 consecutive days at the University of California, Santa Barbara. Sessions occurred once or twice per day (mornings, evenings, or both), yielding 40 MRI sessions, each preceded by state-dependent mood and lifestyle questionnaires and saliva/blood endocrine sampling. At each session, the participant completed a 15-minute eyes-open resting-state fMRI scan on a Siemens 3T Prisma scanner with a 64-channel head coil using a T2*-weighted multiband EPI sequence (TR = 720 ms; TE = 37 ms; flip angle = 56°; 2 mm isotropic voxels; 72 oblique slices; multiband factor = 8), following a high-resolution T1-weighted structural scan and field map. To minimize head motion, the participant’s head was stabilized using a custom 3D-printed foam headcase (CaseForge). A total of 600 minutes (10 hours) of resting-state fMRI data was available for this individual. Raw fMRI data were obtained and preprocessed using the main-text pipeline adapted for single-echo acquisitions, with subsequent denoising using an aCompCor ^83^ framework. The denoised fMRI time-series was mapped to the individual’s fsLR 32k midthickness surfaces with native cortical geometry preserved (using the “-ribbon-constrained” method), combined into the CIFTI format using Connectome Workbench command line utilities^53^.

